# NR2F2 Reactivation in Early-life Adipocyte Stem-like Cells Rescues Adipocyte Mitochondrial Oxidation

**DOI:** 10.1101/2024.09.09.611047

**Authors:** Snehasis Das, Rohan Varshney, Jacob W. Farriester, Gertrude Kyere-Davies, Alexandrea E. Martinez, Kaitlyn Hill, Michael Kinter, Gregory P. Mullen, Prabhakara R. Nagareddy, Michael C. Rudolph

## Abstract

In humans, perinatal exposure to an elevated omega-6 (n6) relative to omega-3 (n3) Fatty Acid (FA) ratio is associated with the likelihood of childhood obesity. In mice, we show perinatal exposure to excessive n6-FA programs neonatal Adipocyte Stem-like cells (ASCs) to differentiate into adipocytes with lower mitochondrial nutrient oxidation and a propensity for nutrient storage. Omega-6 FA exposure reduced fatty acid oxidation (FAO) capacity, coinciding with impaired induction of beige adipocyte regulatory factors PPARγ, PGC1α, PRDM16, and UCP1. ASCs from n6-FA exposed pups formed adipocytes with increased lipogenic genes in vitro, consistent with an in vivo accelerated adipocyte hypertrophy, greater triacylglyceride accumulation, and increased % body fat. Conversely, n6-FA exposed pups had impaired whole animal ^13^C-palmitate oxidation. The metabolic nuclear receptor, NR2F2, was suppressed in ASCs by excess n6-FA intake preceding adipogenesis. ASC deletion of NR2F2, prior to adipogenesis, mimicked the reduced FAO capacity observed in ASCs from n6-FA exposed pups, suggesting that NR2F2 is required in ASCs for robust beige regulator expression and downstream nutrient oxidation in adipocytes. Transiently re-activating NR2F2 with ligand prior to differentiation in ASCs from n6-FA exposed pups, restored their FAO capacity as adipocytes by increasing the PPARγ-PGC1α axis, mitochondrial FA transporter CPT1A, ATP5 family synthases, and NDUF family Complex I proteins. Our findings suggest that excessive n6-FA exposure early in life dampens an NR2F2-mediated induction of beige adipocyte regulators, resulting in metabolic programming that is shifted towards nutrient storage.

## INTRODUCTION

The prevalence of obesity in children and adolescents in the United States is approaching 20%, and approximately 40% of U.S. children are either overweight or obese^1^. Predictions suggest that half of American children may reach clinical obesity by age 35^2^, underscoring the urgency of this public health crisis. The onset of obesity is not only occurring at younger ages but is frequently accompanied by a range of comorbidities typically associated with adulthood, such as non-alcoholic fatty liver, type 2 diabetes, cardiovascular disease, psycho-social consequences, and even premature mortality^3–5^. Although obesity has a complex etiology, including genetic, epigenetic, environmental, and behavioral characteristics, emerging evidence points to early-life nutrition as central to obesity risk, not merely as substrates and fuels for fetal and postnatal development, but as programming cues that can establish enduring consequences for adult metabolism^6–10^. For example, maternal intake of a Western-style diet, rich in omega-6 fatty acids (n6-FA) and processed sugars, has been implicated in metabolic programming changes in offspring, potentially increasing risks for non-communicable metabolic diseases^3^. In recent years, the balance of dietary lipids consumed by pregnant and lactating mothers has come into focus, particularly the ratio of pro-adipogenic n6-FA relative to anti-adipogenic omega-3 (n3) FA ^11–14^.

Maternal transmission of a high ratio of n6/n3 FA to offspring is recognized for its influence in shaping adipocyte development and adipose tissue accumulation^11,12,15–18^. Our research, and that of others, shows that a high n6/n3 FA ratio in breast milk accelerates infant body fat deposition and increases the likelihood of greater childhood adiposity^18–24^. For example, in a cohort of 48 exclusively breastfeeding mother infant dyads, we showed an elevated n6/n3 FA ratio in the milk predicted the change in infant body fat accumulation in the first four months of life, controlling for maternal prepregnancy BMI, fish oil supplementation, gestational weight gain, infant sex, and breastfeeding exclusivity^25^. In neonates particularly, adipose tissue is not merely a nutrient-storage reservoir of lipid-laden adipocytes^26^, it is a critical metabolic and signaling organ that contributes to thermogenesis, through nutrient oxidation, to defend the body temperature of offspring^27–29^. Consistent with our human infant findings, we reported that mouse pups exposed to n6-FA during the fetal and postnatal window had greater body fat at postnatal day (PND) 14, and an inguinal subcutaneous adipose tissue (SAT) with larger, unilocular adipocytes, indicating a nutrient storage phenotype^30^. Molecularly, a high n6/n3 FA ratio exposure increased master adipogenic regulator Peroxisome Proliferator Activated Receptor gamma (PPARγ), lipogenic enzyme mRNA and protein levels, and produced an activated mRNA signature for the “adipogenesis pathway” ^30^.

Adipocyte Stem-like Cells (ASCs) commit to become determined preadipocytes, and then terminally differentiate, giving rise to functional adipocytes^31–35^. More recently, we investigated whether a high n6/n3 FA ratio exposure, specifically during postnatal development, affected adipogenesis by patterning the molecular signature of Adipocyte Stem-like Cells (ASCs)^36^. Through bulk and single-cell RNA-sequencing analyses, we found ASCs from high n6/n3 FA ratio exposed pups had an inhibited b-Catenin (CTNNb1) and Wingless-type MMTV integration site (WNT) mRNA signature, and significantly decreased mRNA and protein levels of Nuclear Receptor 2 group F member 2 (NR2F2, also known as COUP-TF2) ^36^. NR2F2 is a ligand-activated transcription factor that is known to be induced by activation of the WNT/CTNNb1 signaling pathway in 3T3L1 preadipocyte cells^37,38^. NR2F2 plays a pivotal role in regulating metabolism across various tissues, by either activating or repressing transcription through direct DNA binding or dimerization with other nuclear receptors ^39–44^. For example, NR2F2 interaction with PPARα regulates lipid metabolism in the liver, where NR2F2-PPARa cooperativity mediates expression of fatty acid oxidation (FAO) target genes ^45^. Others showed that NR2F2 can bind PPARg, or sequester binding partners such as RXR away from PPARg^46^. We reported distinct ASC responses to *in vivo* n6-FA exposure, in that isolated inguinal SAT ASCs had altered mitochondrial gene expression patterns, fewer “mitochondrial-high” ASCs, and reduced FAO prior to differentiation, which coincided with increased fat mass and larger, unilocular nutrient-storing adipocytes^36^.

Given these early-life high n6/n3 FA ratio effects on inguinal ASCs, we hypothesize that fetal and postnatal exposure to high n6-FA levels trigger ASCs to differentiate into nutrient-storing rather than nutrient-oxidizing adipocytes, which might be overcome by activating NR2F2. We provided pregnant and lactating dams with a specialized diet rich in n6-FA relative to dams provided a balanced, control-FA specialized diet to test the molecular signatures, adipocyte cellular fuel utilization, whole-body metabolic responses, and whether activation of NR2F2 could rescue adipocyte metabolism in ASCs programmed in vivo by n6-FA exposure in PND12 pups. Our findings indicate that in undifferentiated ASCs, NR2F2 acts upstream to establish key regulators of nutrient-oxidizing adipocytes. Furthermore, transient activation of NR2F2 in ASCs isolated from n6-FA exposed pups reignited nutrient-oxidizing metabolism, in part, by restoring mitochondrial protein levels. Our data suggest a model whereby early-life n6-FA exposure limits WNT/CTNNb1 activation, diminishing NR2F2 abundance and robust induction of metabolic regulators, resulting in formation of nutrient-storing adipocytes.

## MATERIALS AND METHODS

### Mouse study

Animal procedures were approved by the IACUC at the University of Oklahoma Health Sciences Center. Wildtype C57BL/6J mice were purchased from Jackson Laboratories (Bar Harbor, MN, USA). The study design used wildtype mothers to assess the effect of different PUFA ratio exposures on offspring development. Prospective dams were provided with the control diet until mating, during which they were divided into control (balanced n6/n3 ratio) and n6-FA (high n6/n3 ratio) groups, provided with appropriate diets, and underwent normal gestation and lactation. At PND12, litters were assessed for body composition, and pups were sacrificed for histology, collection of stromal vascular fractions (SVF), ASC flow cytometry, and assessment of gene expression, protein levels, and circulating hormones, glucose, and fatty acid composition.

### Body composition and indirect calorimetry

Body composition was assessed using quantitative magnetic resonance (qMR; Echo MRI Whole Body Composition Analyzer 4in1-500; Echo Medical Systems, Houston, TX, USA) on PND 12 for dams and litters, as previously described ^30^. Individual litters were weighed before undergoing three consecutive body composition scans, and averages of returned values for fat and lean mass were utilized for analysis.

Dam-litter dyads were placed in indirect calorimetry (IDC, Sable Systems International, Las Vegas, Nevada) cages housed inside the environmental cabinet maintained at 26°C at PND 10. On PND 11, dams were separated from their respective litters for 2 hours, placed back with their litters for 1 hour, then removed from their litters again for 3 hours before finally being placed back with their litters until PND 12. The litters were weighed and body composition measured before the first and second separation of the dam using a weigh-suckle-weigh paradigm to estimate the amount of milk consumed by the pups during the 1 hour feeding window. During the period from PND 10 to PND 12, the dyads (and isolated litters intermittently during weigh-suckle-weigh) in the IDC were monitored for metabolic parameters, including respiratory exchange ratio (RER).

In separate experiments, independent dam-litter dyads were placed in IDC chambers at PND 10. At PND 11, litters were gavaged with 500μg (50µL) of uniformly (^13^C_16_) labeled palmitate (dissolved in peanut oil). The dam was separated from the pups for four hours, during which litter respirometry was measured in the IDC to analyze the amount of labeled (^13^C) CO_2_ exhaled by the litter per minute using a stable isotope analyzer (Sable Systems International). This gave a measure of palmitate oxidation by the litters. Litters were euthanized on PND 12, and tissues collected to measure the ^13^C_16_ palmitate uptake by GC-MS.

### Histology and Immunohistochemistry (IHC)

Histological blocks were prepared from Subcutaneous White Adipose Tissue (SAT). Inguinal SAT pads were gathered from perinatal (PND 10–13) mice, fixed in 10% neutral buffered formalin (NBF), rinsed, and stored in 70% EtOH, and embedded in paraffin. Slides mounted with 5–10 µm sections were analyzed using Immunohistochemistry (IHC) and Hematoxylin and Eosin (H&E) staining. Brightfield and Immunofluorescence whole-slide imaging was conducted using a ZEISS Axio Scan. Z1 Slide Scanner, and composite images were compiled using Zeiss’s Zen Blue software.

Slides used for Hematoxylin and eosin (H&E) staining were prepared following the standard Leica Biosystems’ “Best Practices” protocol. Quantification of adipose cellularity was accomplished through digital analysis of scanned histological sections using the Adiposoft plugin^47,48^ for ImageJ/Fiji^49^. Brightfield whole-tissue scans of H&E-stained slides were optimized for ImageJ processing—including sharpening/clarification and masking of nontarget tissues to reduce artifacts—using Adobe Photoshop. Adipocyte cellularity (diameter, µm) was quantified from each optimized section with Adiposoft. After manually removing any remaining Adiposoft artifacts from the output data, diameter values within a set threshold (15–200 µm) were binned and imported into GraphPad PRISM, where the percentage of adipocytes within each bin was analyzed.

Slides for IHC staining were processed following a standard multi-color immunofluorescence staining protocol^50,51^ and counterstained with DAPI (200 ng/mL). For each sample, multi-channel, shading-corrected images rendered by ZEN Blue were exported into Adobe Photoshop for preparation for image analysis. The tissue sample was isolated from the composite scan by masking background space, nontarget tissues, and instances of highlight clipping, using DAPI and Perilipin-1-stained area for reference when demarking tissue boundaries. Final images used for quantitative analysis were rendered by applying this mask to each color channel image. Additional images were rendered from masked composite scans by creating separate masks that visually isolated beige and white adipose tissue, allowing relative intensity to be quantified independently. Using ImageJ, relative intensity of each fluorophore was determined by finding the quotient of the total area of the tissue sample and the integrated density (the product of the area and the mean grey value, or average gray value of pixels, within a selection) of fluorescence.

Primary antibodies used on SAT sections for IHC were UCP1 (1:200) and Rabbit Perilipin-1 mAb (1:500). Secondary Antibodies utilized were Alexa Fluor 594 Anti-Rat IgG, Alexa Fluor 594 Anti-Rabbit IgG, and Alexa Fluor 488 Anti-Rabbit IgG. Alexa Fluor 594 was analyzed for relative intensity quantification. Adipocytes within white and beige tissue were quantified by exporting the DAPI-Channel image with phenotype masks applied into ImageJ and analyzing using the built-in “Analyze Particles” function. From this, cell density (nuclei/µm^2^) was found by dividing the nuclei count by the area of the target tissue.

### Flow cytometry sorting and analysis

Flow sorting and analyses were performed as previously described^36^. Briefly, subcutaneous adipose tissue was minced and digested in Hanks Balanced Salt Solution (HBSS) (Sigma, H8264) containing 3% BSA, 0.8 mg ml^-1^ collagenase type 2 (Worthington Biochemical, LS004174), 0.8 mM ZnCl_2_, 2.5 mM glucose and 0.2 μM adenosine for 45 minutes at 37°C in an orbital shaker at 150 RPM, and samples were shaken vigorously by hand for one minute after digestion. The resulting suspension was filtered through a 70µm filter. Cells in the stromal vasculature fraction were pelleted at 300xg, washed with HBSS buffer containing 3% BSA, and stained with primary antibodies on ice for 30 minutes. The following antibodies were used: CD45 FITC at 1:1,000 (BioLegend; 103108), CD31 PE-Cy7 at 1:500 (BioLegend, 102417), CD29 Alexa Fluor 700 at 1:200 (BioLegend, 102218), CD34 APC at 1:50 (BioLegend, 119310), Sca-1 BV510 at 1:200 (BioLegend, 108129), and CD24 PE at 1:100 (BioLegend, 138504). Following antibody incubation, cells were washed, and unfixed cell preparations were treated with DAPI (Invitrogen) at 1µg/ml to exclude dead cells. Cells were sorted using a FACS Aria Fusion equipped with FACS DiVA software (BD Biosciences) using a specific gating strategy (Suppl Fig 2). Cell populations were selected based on forward scatter (FSC) and side scatter (SSC), and dead cells were excluded. Single cells were isolated or analyzed based on cell surface markers. Data was analyzed using BD FACS DiVA and FlowJo.

### Cell culture and differentiation

The immortalized adipocyte precursor (mAPC) cell line was purchased from Kerafest (MA, USA). mAPCs were cultured in High Glucose Dulbecco’s Modified Eagle Medium 1:1 with Hams F-12 (DMEM/F12) containing 10% FBS supplemented with penicillin (100 U/mL) and streptomycin (100 µg/mL) at 37℃ in a humidified incubator with 5% CO_2_. Primary ASCs were cultured under similar conditions in High Glucose Dulbecco’s Modified Eagle Medium (DMEM) containing 10% fetal bovine serum (FBS) supplemented with penicillin and streptomycin. For differentiating cells into beige adipocytes, mAPCs and ASCs were cultured in their respective mediums until nearly confluent and the medium was then changed to induction medium (5 μg/mL insulin, 1 nM T3, 125 μM indomethacin, 2 μg/mL dexamethasone, 0.5 mM IBMX, 0.5 μM rosiglitazone) when cells are 80-90% confluence (day 2). After 48h (Day 4), the medium was changed to maintenance medium (5 μg/mL insulin, 1 nM T3) with 0.5 μM rosiglitazone). On day 6 and 8, spent medium was replaced with fresh maintenance medium with 1 μM rosiglitazone. On termination of differentiation at day 10, cells were utilized for the various assays described herein. For some experiments, 1-DSO (300 nM) was added to ASCs and mAPCs for four days flanking induction (two days before the induction to two days after). This is depicted schematically in the figures.

For NR2F2 knockdown experiments, ASCs isolated from NR2F2^f/f^ mice were transduced with adeno-Cre (AdCre) or adeno control (AdCon) virus. 5-6 days after transduction, the cells were treated with induction media to initiate differentiation. For other experiments, 70-80% confluent ASCs from n6-FA pups were treated with Wnt agonist 1 for 48h to induce Wnt-CTNNb1 pathway.

### Live cell staining for mitochondrial potential, fatty acid oxidation, and lipid accumulation

Primary ASCs were cultured as described above. For staining with tetramethylrhodamine ethyl ester (TMRE, Biotium) live cell mitochondrial potential dye and LipidSpot™ 488 (Biotium), media were changed to fresh DMEM/10% FBS containing these stains diluted 1:1,000. Cells were incubated at 37℃ for 15 minutes before fluorescent data was collected using a Zoe^TM^ Fluorescent Cell Imager (BioRad). For staining with FAOBlue (Funakoshi), TMRE, and LipidSpot 488, cells were washed twice in serum-free DMEM and then incubated for 15 min. at 37℃ in fresh DMEM (no serum) containing FAOBlue (1:1,000). TMRE and LipidSpot™ 488 were added and cells incubated 15 min. at 37℃ before imaging.

### Quantitative PCR and Western blotting

Total RNA was extracted using the RNeasy Plus Mini Kit (Qiagen, Hilden, Germany, 74134) according to the manufacturer’s protocol, and 500ng of total RNA was reverse transcribed into cDNA using iScript Reverse Transcription Supermix (Bio-Rad, Hercules, CA, USA) or the Verso cDNA Synthesis Kit (ThermoFisher, MA, USA). cDNA representing 25ng of total RNA was added to each qPCR reaction containing TaqMan Fast Advanced Master Mix and primers (ThermoFisher, MA, USA) specific for Pparγ, Pgc1α, Ucp1, PRDM16, Cidea, Cox8b, Cox7a1, Acly, Acc1, Fasn, Srebp1, Srebp2, Dgat1, or Nr2f2 were used for quantitative real time PCR (Applied Biosystem, 589 Ca, USA). Ef1α was used as a housekeeping gene. mRNA levels were determined using the ΔΔCT method and normalized to the housekeeping gene Ef1α.

For Simple Westerns using JESS, flow-sorted primary ASCs, mature adipocytes or whole iWAT were homogenized in RIPA buffer with protease and phosphatase inhibitors. Lysates were run on a ProteinSimple JESS instrument for PPARγ (CST-2435), PGC1α (CST-2178), C/EBPα (CST-2295), UCP1 (CST-14670), CPT1A (Protein Tech-15184-1-AP), CPT1C (Protein Tech-12969-1-AP), FABP4 (CST-2120), NR2F2 (CST-6434), SOD2 (CST-13141), TOM20 (CST-42406), and PLIN1 (CST-9349) antibodies. All antibodies were diluted 1:50 and 3 μL of 1.2mg/ml protein/well was loaded for JESS Westerns. Protein from primary ASCs from six different SVFs for each dietary condition (high n6 or control) were used for each JESS assay.

### Seahorse Cellular Metabolic Assays

Seahorse substrate oxidation assays were conducted using the manufacturer’s instructions with some modifications described below, utilizing the glucose/pyruvate oxidation stress test kit (103673-100), glutamine oxidation stress test kit (103674-100), and palmitate oxidation stress test kit (103693-100). Modifications include adding inhibitors (20µL per well) directly to the cells both 15 minutes before the assay and following the standard mito stress test protocol.

### Targeted Lipid Analysis

500µL of 0.1M potassium phosphate buffer, pH 6.8 was added to 13×100mm borosilicate glass culture tubes (Fisher Scientific, #14-961-27), to which 6µL of pup serum was added. The mixture was then acidified with 10µL of 1N HCl (prewashed with Hexanes) and vortexed briefly. 500µL of methanol was added to this solution and vortexed briefly. Total lipids were extracted twice by adding 1mL of isooctane:ethyl acetate 3:1 (vol/vol) and once with 1mL Hexane; for each extraction, samples were vortexed vigorously for 10-15 sec and centrifuged at 2000g for 1 minute to complete phase separation. The organic layer from each extraction was combined into a new 13×100 mm borosilicate glass tube. Extracted lipids were brought to dryness and resuspend in 300µL 2,2,4-Trimethylpentane (isooctane). For the non-esterified fatty acid (NEFA) fraction, 100µL of resuspended total lipids was transferred to a new borosilicate glass tube, mixed with (25 ng) blended stable isotope internal standard, taken to dryness under gaseous N_2_, and resuspended in 25µL of 1% pentafluorobenzyl bromide in acetonitrile (vol/vol), to which 25µL of 1% diisopropylethylamine in acetonitrile (vol/vol) was added, and samples were incubated at room temperature for 30 minutes. Pentafluorobenzyl-fatty acid derivatives were taken to dryness under gaseous N_2_ and resuspended in 100µL hexane for injection into the GC/MS. For the total fatty acid fraction (TFA), 50µL of the original 300µL total lipid extract was transferred to a separate Teflon lined screw cap glass tube, mixed with (66.7 ng) blended stable isotope internal standard, and taken to dryness under gaseous N_2_. TFA samples were resuspended in 500µL of 100% ethanol, to which 500µL of 1M NaOH was added to saponify the TFA fraction at 90°C for 30 minutes, followed by acidification using 550µL of 1M HCl. Saponified samples were then extracted twice with 1.5mL of Hexanes, taken to dryness under gaseous N_2_, and derivatized as above. Derivatized TFA samples were resuspended in 267µL hexanes for injection into the GC/MS. For ^13^C_16_-palmitate measurement in tissues from tracer gavaged pups, the tissues were homogenized in a bead mill (Fisher Scientific) in 2mL bead mill tubes in 1mL of 66% methanol (in pH 6.8 KPhos buffer). The homogenates were transferred to 13×100mm borosilicate glass tubes and 40µL of prewashed HCl was added mixed briefly. Lipids were extracted as above and reconstituted in 300µL Isooctane. 50µL of resuspended total lipids was transferred to a new Teflon lined screw cap glass tube, mixed with (66.7 ng) ^2^D_31_-palmitate internal standard and taken to dryness under gaseous N_2_. Samples were saponified, extracted, and derivatized as above. Derivatized samples were resuspended in 267µL hexanes for injection. For the NEFA, TFA and tracer fractions, 1µL of pentafluorobenzyl-fatty acid derivatives was injected and data were collected on the GC-MS (8890 GC, 5977B MSD, Agilent) DB-1MS UI column (122-0112UI, Agilent) with the following run program: 80°C hold for 3 minutes, 30°C/minute ramp to 125°C, no hold, 25°C ramp to 320°C and hold for 2 minutes. The flow rate for the methane carrier gas was set at 1.5mL/minute. Data were acquired in full scan negative chemical ionization mode to identify fatty acids of acyl chain length from 8 to 22 carbons. Peak areas of the analyte or standard were measured, and the ratio of the area from the analyte-derived ion to that from the internal standard was calculated^52^. ^13^C_16_-palmitate was analyzed relative to the quantitative ^2^D_31_-palmitate internal standard^53^.

### Proteomics

ASCs were isolated from control and n6-FA pups and plated in 6-well plates, followed by differentiation into beige adipocytes, with or without 1-DSO exposure flanking beige induction in the n6-FA ASCs. After differentiation, cells were washed twice with PBS and lysed in 100ml RIPA buffer per well and used for proteomics.

### Sample processing for proteomics

A GeLC approach was used, in which each sample was run approximately four cm into an SDS-Page gel, fixed, and stained with Coomassie blue. Each lane of the gel was cut top to bottom as a series of seven broad molecular weight fractions. ach fraction was chopped into smaller pieces, washed, reduced with DTT, alkylated with iodoacetamide, and digested with 1µg trypsin overnight at room temperature. Peptides were extracted from the gel in 50% acetonitrile, the extracts taken to dryness by Speedvac, and reconstituted in 200µL 1% acetic acid for analysis.

### LC-MS for proteomics

We used a ThermoScientific QEx Plus instrument interfaced to an Ultimate 3000 nanoflow HPLC system with 75um x 20xm C18 (Phenomenex Aeris XB-18 3.6um beads) capillary columns packed in CoAnn tips. 10uL samples are injected and loaded on the column at 1.5uL/min for 8minutes. Peptides are eluted with a linear gradient of acetonitrile in water with 0.1% formic acid from 1% to 50% in 60min at a flow rate of 150nL/min.

### Data dependent acquisition (DDA)

The DDA experiments used a top 20 strategy where one full scan MS spectrum was acquired followed by 20 CID spectra. The full scan mass spectra were acquired with an m/z resolution of 70,000 and CID spectra with an m/z resolution of 17,500. Ion source settings included a spray voltage of 2.2 kV, ion transfer tube temperature of 300◦ C, and positive ions mode.

DDA data are searched against the Uniprot proteome database from Embl. Mascot matches each CID spectrum to a peptide sequence in the database, considering the digestion with trypsin, cysteine alkylation, and variable methionine oxidation. The identifications are ranked based on a total score determined by the program based on the quality and number of peptide matches. Only proteins with two or greater matching peptides are included in the results.

### Enzyme linked immunosorbent assay (ELISA)

ELISAs for high molecular weight (HMW) and total adiponectin (ALPCO, 47-ADPMS-E01), Acylation Stimulating Protein (ASP, MyBioSource, MBS263213), Complement C3 (Abcam, ab157711), Insulin (Crystal Chem, 90080), Leptin (R&D, DY498) and Retinol-Binding Protein 4 (RBP4, R&D, DY3476) were performed following manufacturers’ instructions. Pup serum was collected from trunk blood (pooled from two to three pups per group) and diluted 1:8181, 1:200, 1:50000, 1:1, 1:1 and 1:2000 for HMW and total adiponectin, ASP, Complement C3, Insulin, Leptin and RBP4, respectively.

### Statistical Analyses

Results are presented as meanJ±JSEM with *p* values less than 0.05 considered significantly different. Two-way ANOVA, repeated measures two-way ANOVA, one-way ANOVA, t-test, or multiple comparison t-test were performed for the analyses of differences between groups. Tukey post-hoc comparisons were done where appropriate using GraphPad Prism Software, Version 9.

## DATA AVAILABILITY

Source data are provided with this paper. Source data for Figures 4L and 5H are included with this paper as source data file 1. Source data for Table 1 is included in this paper as Source data file 2. No publicly available data sets were generated or analyzed for this study.

**Table 1:**
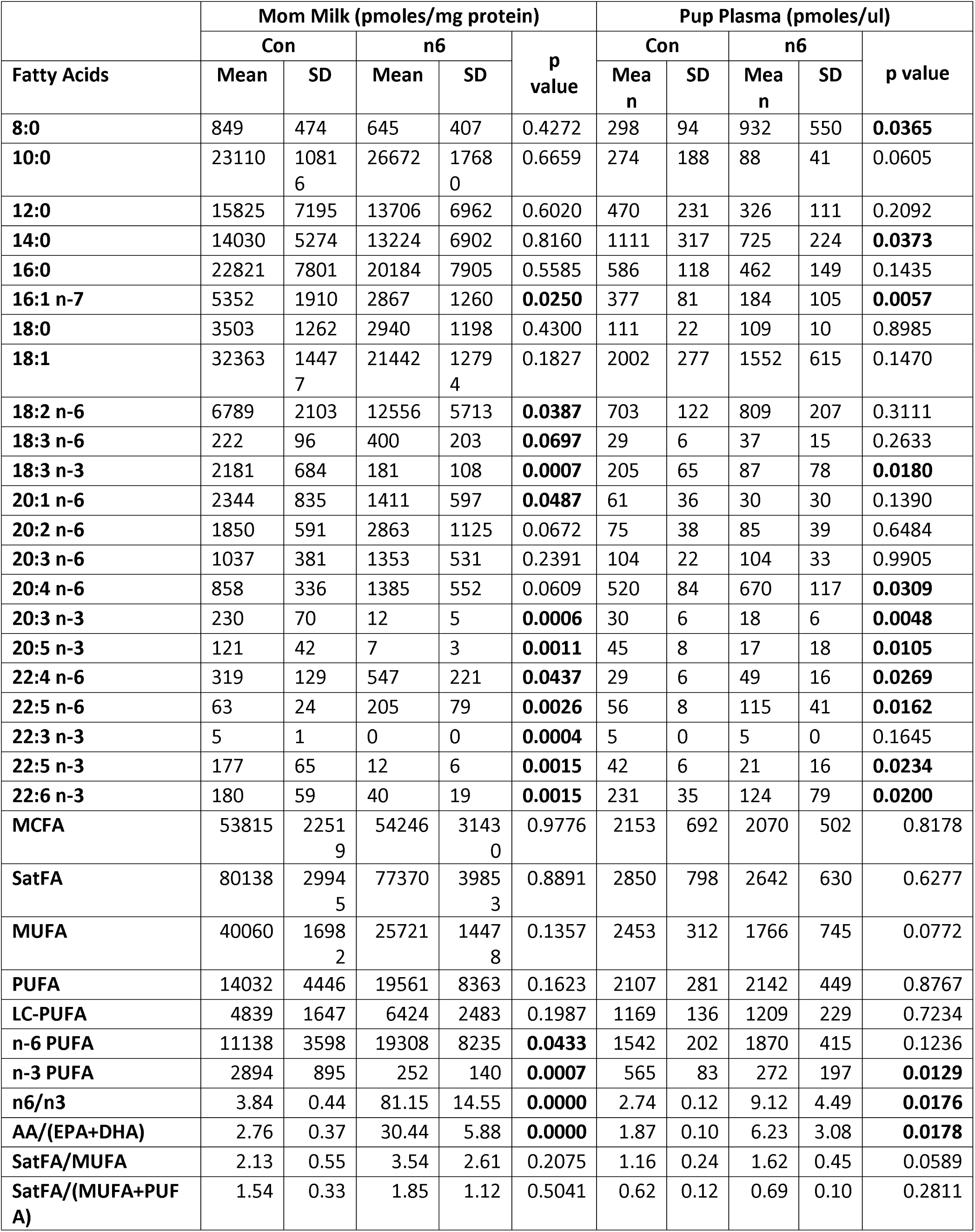
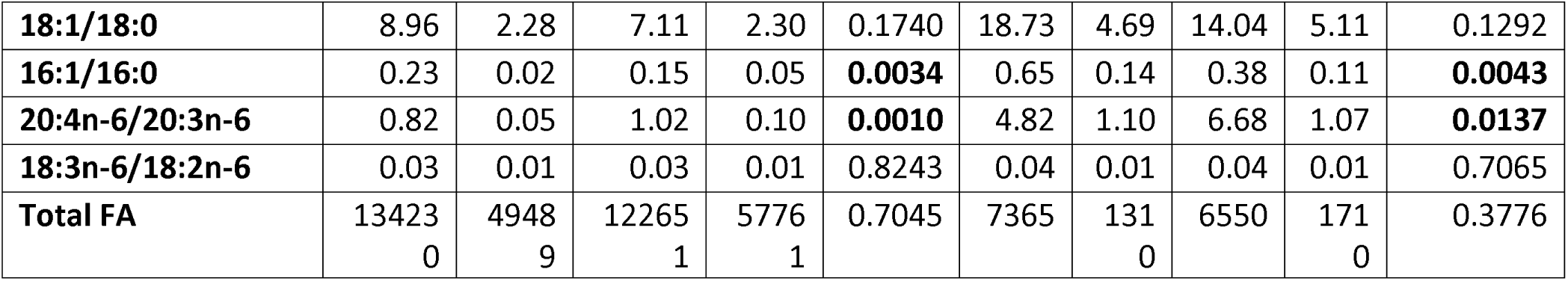
GC-MS analyses of mother’s milk and plasma from the pups for quantifying levels of omega-3 (n3) and omega-6 (n6) fatty acids (n=5-7).

|  | Mom Milk (pmoles/mg protein) |  |  |  |  | Pup Plasma (pmoles/ul) |  |  |  |  |
| --- | --- | --- | --- | --- | --- | --- | --- | --- | --- | --- |
|  | Con |  | n6 |  | p value | Con |  | n6 |  | p value |
| Fatty Acids | Mean | SD | Mean | SD |  | Mean | SD | Mean | SD |  |
| 8:0 | 849 | 474 | 645 | 407 | 0.4272 | 298 | 94 | 932 | 550 | **0.0365** |
| 10:0 | 23110 | 10816 | 26672 | 17680 | 0.6659 | 274 | 188 | 88 | 41 | 0.0605 |
| 12:0 | 15825 | 7195 | 13706 | 6962 | 0.6020 | 470 | 231 | 326 | 111 | 0.2092 |
| 14:0 | 14030 | 5274 | 13224 | 6902 | 0.8160 | 1111 | 317 | 725 | 224 | **0.0373** |
| 16:0 | 22821 | 7801 | 20184 | 7905 | 0.5585 | 586 | 118 | 462 | 149 | 0.1435 |
| 16:1 n-7 | 5352 | 1910 | 2867 | 1260 | **0.0250** | 377 | 81 | 184 | 105 | **0.0057** |
| 18:0 | 3503 | 1262 | 2940 | 1198 | 0.4300 | 111 | 22 | 109 | 10 | 0.8985 |
| 18:1 | 32363 | 14477 | 21442 | 12794 | 0.1827 | 2002 | 277 | 1552 | 615 | 0.1470 |
| 18:2 n-6 | 6789 | 2103 | 12556 | 5713 | **0.0387** | 703 | 122 | 809 | 207 | 0.3111 |
| 18:3 n-6 | 222 | 96 | 400 | 203 | **0.0697** | 29 | 6 | 37 | 15 | 0.2633 |
| 18:3 n-3 | 2181 | 684 | 181 | 108 | **0.0007** | 205 | 65 | 87 | 78 | **0.0180** |
| 20:1 n-6 | 2344 | 835 | 1411 | 597 | **0.0487** | 61 | 36 | 30 | 30 | 0.1390 |
| 20:2 n-6 | 1850 | 591 | 2863 | 1125 | 0.0672 | 75 | 38 | 85 | 39 | 0.6484 |
| 20:3 n-6 | 1037 | 381 | 1353 | 531 | 0.2391 | 104 | 22 | 104 | 33 | 0.9905 |
| 20:4 n-6 | 858 | 336 | 1385 | 552 | 0.0609 | 520 | 84 | 670 | 117 | **0.0309** |
| 20:3 n-3 | 230 | 70 | 12 | 5 | **0.0006** | 30 | 6 | 18 | 6 | **0.0048** |
| 20:5 n-3 | 121 | 42 | 7 | 3 | **0.0011** | 45 | 8 | 17 | 18 | **0.0105** |
| 22:4 n-6 | 319 | 129 | 547 | 221 | **0.0437** | 29 | 6 | 49 | 16 | **0.0269** |
| 22:5 n-6 | 63 | 24 | 205 | 79 | **0.0026** | 56 | 8 | 115 | 41 | **0.0162** |
| 22:3 n-3 | 5 | 1 | 0 | 0 | **0.0004** | 5 | 0 | 5 | 0 | 0.1645 |
| 22:5 n-3 | 177 | 65 | 12 | 6 | **0.0015** | 42 | 6 | 21 | 16 | **0.0234** |
| 22:6 n-3 | 180 | 59 | 40 | 19 | **0.0015** | 231 | 35 | 124 | 79 | **0.0200** |
| MCFA | 53815 | 22519 | 54246 | 31430 | 0.9776 | 2153 | 692 | 2070 | 502 | 0.8178 |
| SatFA | 80138 | 29945 | 77370 | 39853 | 0.8891 | 2850 | 798 | 2642 | 630 | 0.6277 |
| MUFA | 40060 | 16982 | 25721 | 14478 | 0.1357 | 2453 | 312 | 1766 | 745 | 0.0772 |
| PUFA | 14032 | 4446 | 19561 | 8363 | 0.1623 | 2107 | 281 | 2142 | 449 | 0.8767 |
| LC-PUFA | 4839 | 1647 | 6424 | 2483 | 0.1987 | 1169 | 136 | 1209 | 229 | 0.7234 |
| n-6 PUFA | 11138 | 3598 | 19308 | 8235 | **0.0433** | 1542 | 202 | 1870 | 415 | 0.1236 |
| n-3 PUFA | 2894 | 895 | 252 | 140 | **0.0007** | 565 | 83 | 272 | 197 | **0.0129** |
| n6/n3 | 3.84 | 0.44 | 81.15 | 14.55 | **0.0000** | 2.74 | 0.12 | 9.12 | 4.49 | **0.0176** |
| AA/(EPA+DHA) | 2.76 | 0.37 | 30.44 | 5.88 | **0.0000** | 1.87 | 0.10 | 6.23 | 3.08 | **0.0178** |
| SatFA/MUFA | 2.13 | 0.55 | 3.54 | 2.61 | 0.2075 | 1.16 | 0.24 | 1.62 | 0.45 | 0.0589 |
| SatFA/(MUFA+PUFA) | 1.54 | 0.33 | 1.85 | 1.12 | 0.5041 | 0.62 | 0.12 | 0.69 | 0.10 | 0.2811 |
| **18:1/18:0** | 8.96 | 2.28 | 7.11 | 2.30 | 0.1740 | 18.73 | 4.69 | 14.04 | 5.11 | 0.1292 |
| **16:1/16:0** | 0.23 | 0.02 | 0.15 | 0.05 | **0.0034** | 0.65 | 0.14 | 0.38 | 0.11 | **0.0043** |
| **20:4n-6/20:3n-6** | 0.82 | 0.05 | 1.02 | 0.10 | **0.0010** | 4.82 | 1.10 | 6.68 | 1.07 | **0.0137** |
| **18:3n-6/18:2n-6** | 0.03 | 0.01 | 0.03 | 0.01 | 0.8243 | 0.04 | 0.01 | 0.04 | 0.01 | 0.7065 |
| **Total FA** | 13423<br>0 | 4948<br>9 | 12265<br>1 | 5776<br>1 | 0.7045 | 7365 | 131<br>0 | 6550 | 171<br>0 | 0.3776 |

## RESULTS

### Omega-6 FA accelerates fat accumulation and reduces lipid acid oxidation

Female mice were randomized and provided either a control-FA diet based on soy oil or a safflower oil diet rich in n6-FA at the time of mating (Fig 1A). All dams went through normal gestation and parturition, and litters were standardized to 7-8 pups per dam. All mating pairs, dams, and litters were housed in a temperature-controlled satellite facility maintained at 25°C. Gas Chromatography-Mass Spectrometry (GC-MS) analysis of plasma confirmed that n6-FA exposed litters (i.e., offspring of dams consuming diet rich in n6-FA) had significantly higher circulating levels of n6-FA, including arachidonic acid (20:4 n6), adrenic acid (22:4 n6), alongside reduced n3 fatty acids, including linolenic acid (18:3 n3), eicosatrienoic acid (20:3 n3), eicosapentaenoic acid (EPA 20:5 n3), docosapentaenoic acid(20:5 n6), and docosahexaenoic acid (DHA 22:6 n3; Table 1, Source Data File 2).

**Figure 1.**
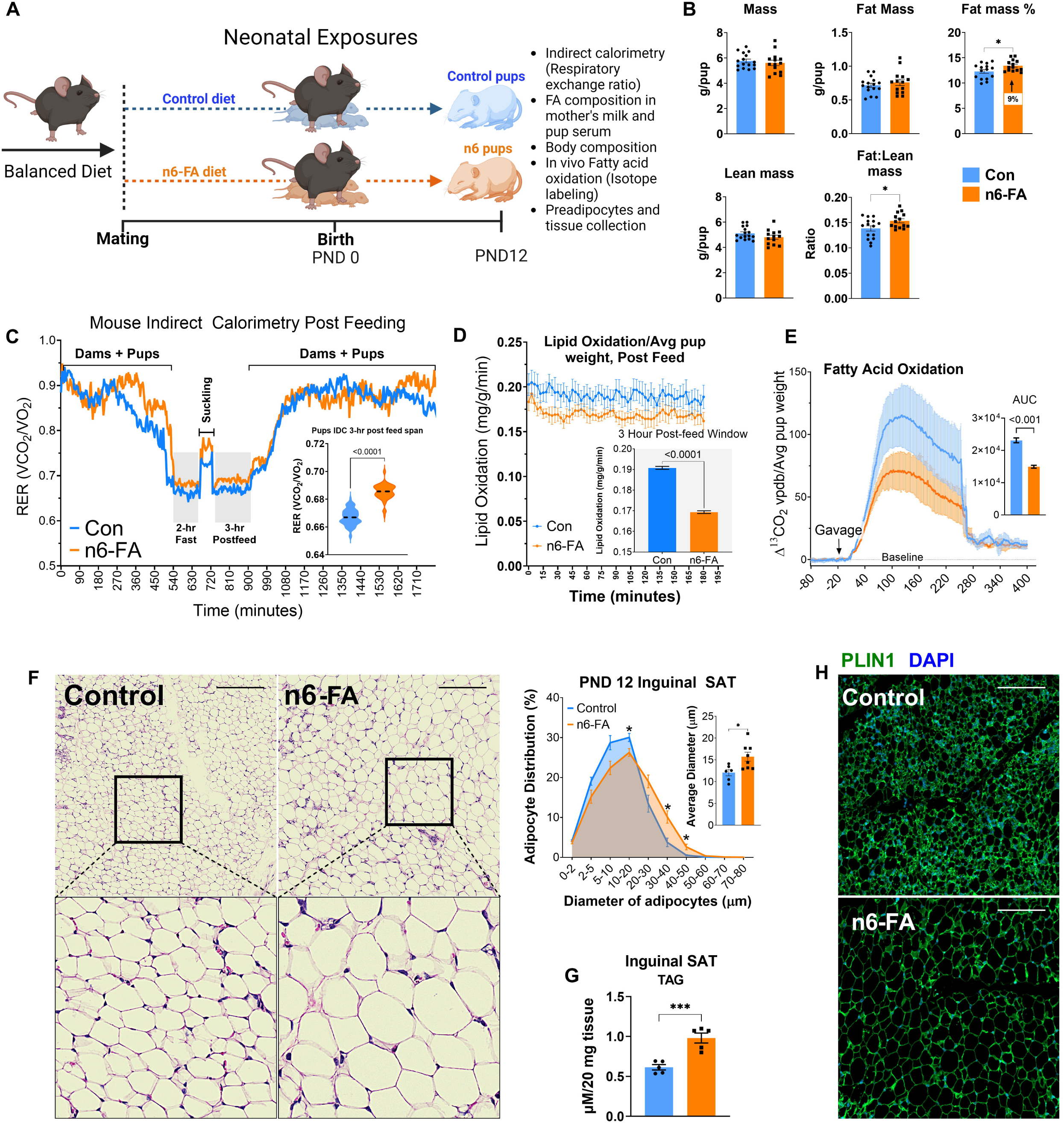
High n6 exposure alters fat mass percentage, RER, and adipocyte cellularity. (A) Diagram of experimental design. (B) Body composition of control and high n6-FA exposed pups at Postnatal day 12 (PND12). Each data point is the mean mass, lean body mass, fat mass, ratio of fat to lean mass, or fat mass percentage of one litter of pups at PND12 (n=13-15 litters). (C) Indirect calorimetry of control and high n6-exposed litters. Dams were removed for the 2h pre- and 3h post-nursing periods to measure the Respiratory Exchange Ratio (VCO2/VO2: Scale 0.5-1.0) of pups in a litter. Data are presented as means ± SEM (n=7-9). (D) Fatty acid oxidation (FAO) calculated using the equation (1.70 × VO_2_ – 1.69 × VCO_2_) (n=7-9). (E) Pups were administered ^13^C_16_-palmitate (100 mg/kg body weight) and the litters placed into cages in an Indirect Calorimeter cabinet held at 25°C to quantify ^13^CO_2_ stable isotope gas exchange as a measure of whole-body FAO (n=3 per group). (F) H&E staining of control and n6-FA exposed pup SAT (inguinal fat) and quantification of cellularity (n=7-8). Scale bar is 100 μm. (G) Quantification of SAT triglyceride (TAG) levels (n=5). (H) Immunofluorescence staining of inguinal adipose tissue with anti-PLIN1 antibody (green) and DAPI (blue) (n=5 neonatal inguinal SAT sections from independent litters per control/n6-FA group). The scale bar is 100 μm. Data are expressed as mean ± SEM, statistical significance is denoted by *p < 0.05, **p < 0.01, ***p < 0.001, ****p < 0.0001 by t-test.

On PND12, body composition analysis by quantitative magnetic resonance (qMR) was assessed prior to pup sacrifice. Litters in the n6-FA exposure group had approximately 9% more body fat (*p*=0.029, n=15 litters/group) and a significantly greater fat-to-lean mass ratio (*p*=0.0322) than control-FA exposed pups, while no differences were observed in overall total body weight or lean mass (Fig 1B). Using indirect calorimetry with the environmental cabinet maintained at 25°C, PND12 litters in the n6-FA group had a significantly higher respiratory exchange ratio (RER, VCO_2_/VO_2_; *p*<0.0001, n=7-9 litters/group) compared to control-FA litters, both before and after suckling (Fig 1C). FAO calculated using the VO_2_ and VCO_2_ data in the post feeding window indicate reduced lipid oxidation in n6-FA litters (Fig 1D).

A separate cohort of n6-FA and control-FA litters were administered ^13^C_16_-palmitate (100 mg/kg body weight) by gavage, placed into calorimetry cages, and whole-body FAO was used to quantify ^13^CO_2_ emission by stable isotope gas exchange. The n6-FA exposed pups had significantly lower ^13^CO_2_ vpdb emission, indicating diminished whole-body FAO (Fig 1E, *p*<0.001, n=4 litters/group). No differences were observed in blood glucose, triacylglycerides, or insulin concentrations between groups, and ^13^C_16_-palmitate uptake, quantified by GC/MS, into inguinal SAT, brown adipose tissue (BAT), liver, and muscle was equivalent (Suppl Fig 1A, B).

### Omega-6 FA pups have less beige fat and reduced UCP1 levels

We investigated the cellular morphology of inguinal SAT using H&E-stained sections. Consistent with our previous findings^36^, n6-FA adipocytes were significantly larger than those from control pups (Fig 1F, *p*<0.05, n=6). Total TAG accumulation in the dissected inguinal SAT depot was significantly increased (*p*=0.021, n=3), consistent with the larger unilocular adipocytes and 9% increased body fat in the n6-FA pups (Fig 1G). No significant changes were observed in serum levels of key adipokines such as high molecular weight and total adiponectin, leptin, and retinol-binding protein 4 (RBP4) (Suppl Fig 1C).

Immunofluorescence for Perilipin1 (PLIN1), a coat protein for cytoplasmic lipid droplets within adipocytes, suggested that multilocular beige adipose tissue (BeAT) in inguinal SAT of n6-FA exposed pups was reduced (Fig 1H). Portions of adipose tissue having multilocular BeAT adipocyte regions, assessed by PLIN1 morphological inspection, were selected to quantify mitochondrial uncoupling protein 1 (UCP1), a marker of nutrient-oxidizing BeAT (Fig 2A). The distribution of UCP1 high staining overlapped with multilocular septa of beige adipocytes in inguinal SAT. Levels of UCP1 in multilocular adipocytes were significantly decreased within BeAT septa (*p*=0.0008, n=9/condition) in n6-SA SAT, followed by a trend of decreased UCP1 in the whole section (*p*=0.052), in the n6-FA sections. We confirmed the significant UCP1 decrease by immunoblotting, using the contralateral inguinal SAT from n6-FA and control-FA pups (Fig 2B, *p*=0.027, n=3/group). The ratio of mitochondrial superoxide dismutase 1 (SOD2) to mitochondrial import receptor subunit TOM20 in adipocytes is an indicator of mitochondrial dysfunction in obesity^54^. By PND12, inguinal SAT of n6-FA exposed pups had a significant increase in the SOD2/TOM20 ratio (Fig 2C, *p*=0.047, n=6/group), despite equivalent mitochondria amounts based on total TOM20 levels.

**Figure 2.**
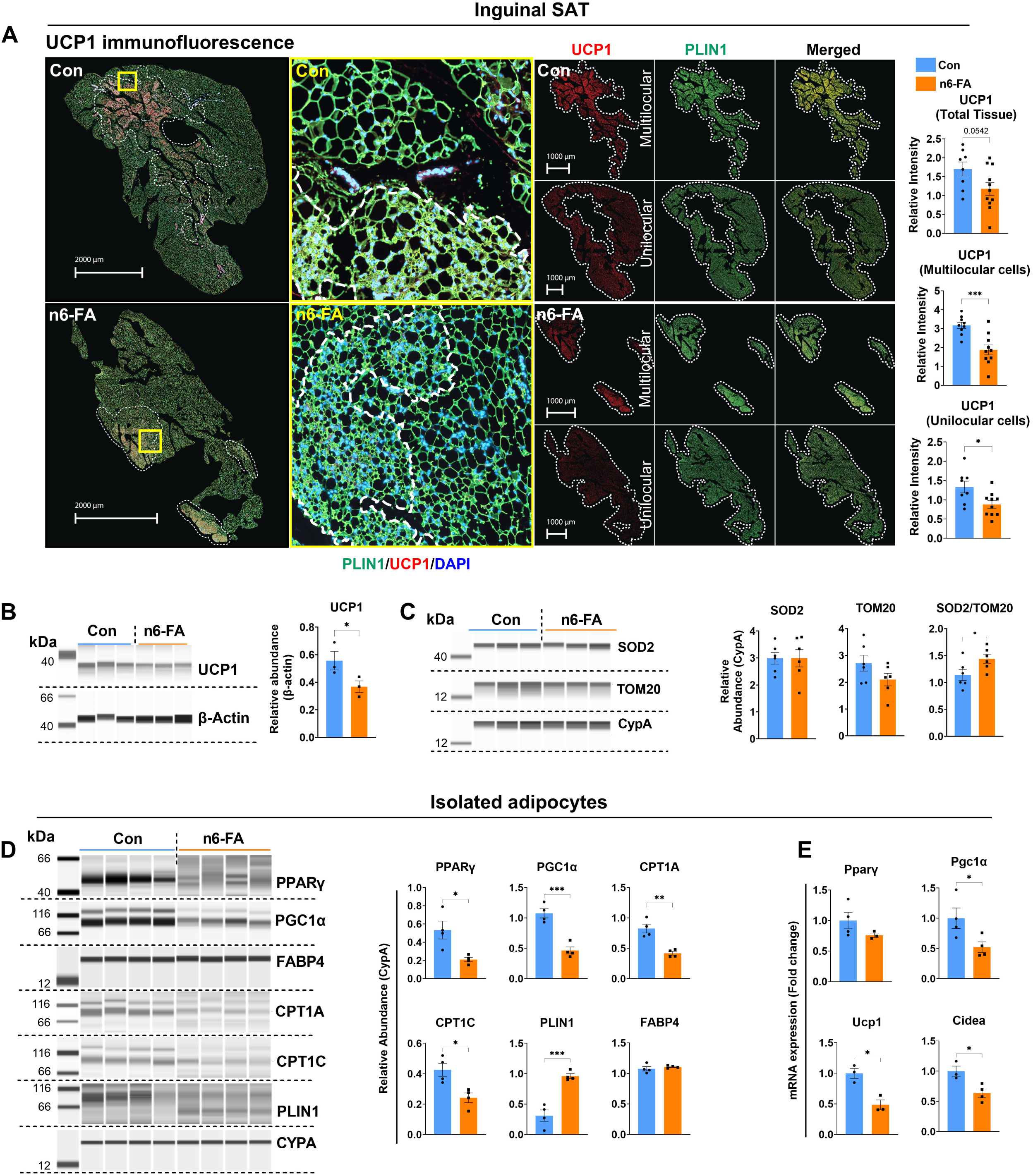
Expression of beige adipocyte proteins in SAT and adipocytes. (A) Immunofluorescence staining of inguinal SAT from PND12 mice with anti-UCP1 (red) and anti-PLIN1 (green) antibodies. Nuclei (blue) were stained with DAPI. Morphologically defined white and beige regions were quantified separately for UCP1 staining, which was significantly decreased in n6-FA group (n=8-11 neonatal SAT section from independent litters per control/n6 group). (B, C) Analysis of UCP1, SOD2, and TOM20 in inguinal fat pads through Western blotting. CypA was used as the reference protein to normalize loading (n=3-6 neonatal SAT tissue from independent litters per control/n6-FA group). (D, E) Levels of key adipocyte proteins and mRNA were measured through Western blotting and qPCR in mature adipocytes isolated from inguinal fat from control and n6-FA pups. (n=3-4 pooled adipocytes from neonatal SAT tissue from independent litters per control/n6-FA group). Data are expressed as mean ± SEM, statistical significance is denoted by *p < 0.05, **p < 0.01, ***p < 0.001, ****p < 0.0001 by t-test.

### Isolated adipocytes of n6-FA SAT have diminished BeAT regulators

We collected mature adipocytes from SVF preparations to assess protein levels and gene expression. The principal regulators of the nutrient-oxidizing adipocyte gene expression program, PPARγ and PPARγ coactivator 1 alpha (PGC1α)^55^, were significantly decreased at the protein level in isolated adipocytes of the n6-FA exposed pups (Fig 2D, *p*=0.018 and 0.0007, respectively, n=4 individual litters/group). Moreover, critical transporters necessary for mitochondrial FAO, carnitine palmitoyl transferase 1A (CPT1A) and CPT1C, were also significantly decreased (*p*=0.002 and 0.013, respectively). Interestingly, the lower band for cytoplasmic lipid droplet protein PLIN1, which coordinates lipase access to the TAG droplet, was sharply increased in isolated adipocytes from n6-FA exposed pups (*p*=0.0007). The expression of genes necessary for nutrient-oxidation in beige adipocytes, Pgc1α, Ucp1, and Cell Death Inducing DFFA Like Effector A (Cidea), was notably decreased (*p*<0.05), while mRNA levels for PPARγ did not differ in isolated adipocytes from n6-rich pups (Fig 2E). Similar gene expression differences were observed in the whole inguinal adipose tissue (Suppl Fig 1D). Given these findings, we investigated whether *in vivo* n6-FA exposure alters the differentiation potential of ASCs by impacting the adipogenic regulators of BeAT.

### Omega-6 FA program ASCs with decreased nutrient-oxidation and increased lipogenesis

We isolated inguinal SAT ASCs for blood and endothelial lineage negative (CD45-, CD31-), mesenchymal stem cell positive (CD29+, CD34+), and adipocyte precursor positive (Sca1+) cells from n6-FA and control-FA exposed PND12 pups by fluorescence activated cell sorting from stromal vascular fraction (SVF) preparations^36^. ASCs were plated, allowed to adhere, and reach confluency, and then differentiated using standard adipogenic hormones, followed by six days in adipocyte maintenance media (Fig 3A). Differentiated adipocytes were stained using live-cell dyes for FAO (FAO blue), mitochondrial activity (TMRE), and triacylglyceride accumulation (LipidSpot). ASCs from the n6-FA exposed inguinal SAT that were differentiated *in vitro* had significantly decreased FAO staining and mitochondrial activity (*p*=0.028 and 0.022, respectively), while overall lipid accumulation was not significantly changed (Fig 3B). Consistent with observations in isolated inguinal SAT adipocytes (Fig 2D and E), following *in vitro* differentiation of isolated ASCs, PPARγ, PGC1α, UCP1, and CPT1A protein levels were significantly decreased by in vivo n6-FA exposure (Fig 3C). Interestingly, no significant difference was observed for C/EBPα, which is needed for induction of PPARγ^56^, and equivalent levels of FABP4 indicated both n6-FA and control ASCs differentiated into adipocytes. Given the sharp reduction in PPARγ and PGC1α protein levels in the n6-rich *in vitro* differentiated adipocytes, we measured gene expression for additional BeAT specific markers. In addition to significantly decreased Pparγ, Pgc1α, and Ucp1, we observed significantly lower expression of Cidea, Prdm16, Cox8b, and Cox7a1, in n6-FA ASCs following differentiation (Fig 3D). In contrast to the beige adipocyte program, the lipogenic markers, ATP-citrate lyase (Acly), Acetyl-CoA Carboxylase (Acc1), and Fatty Acid Synthase (Fasn) were significantly increased, as was expression of Adiponectin (AdipoQ), while Srebp2, the key player for cholesterol biosynthesis tended to increased expression (Fig 3D). Together, these findings suggest that *in vivo* n6-FA exposure programs the adipogenic potential of ASCs for nutrient storage, due to the less robust induction of beige adipocyte gene regulators.

**Figure 3.**
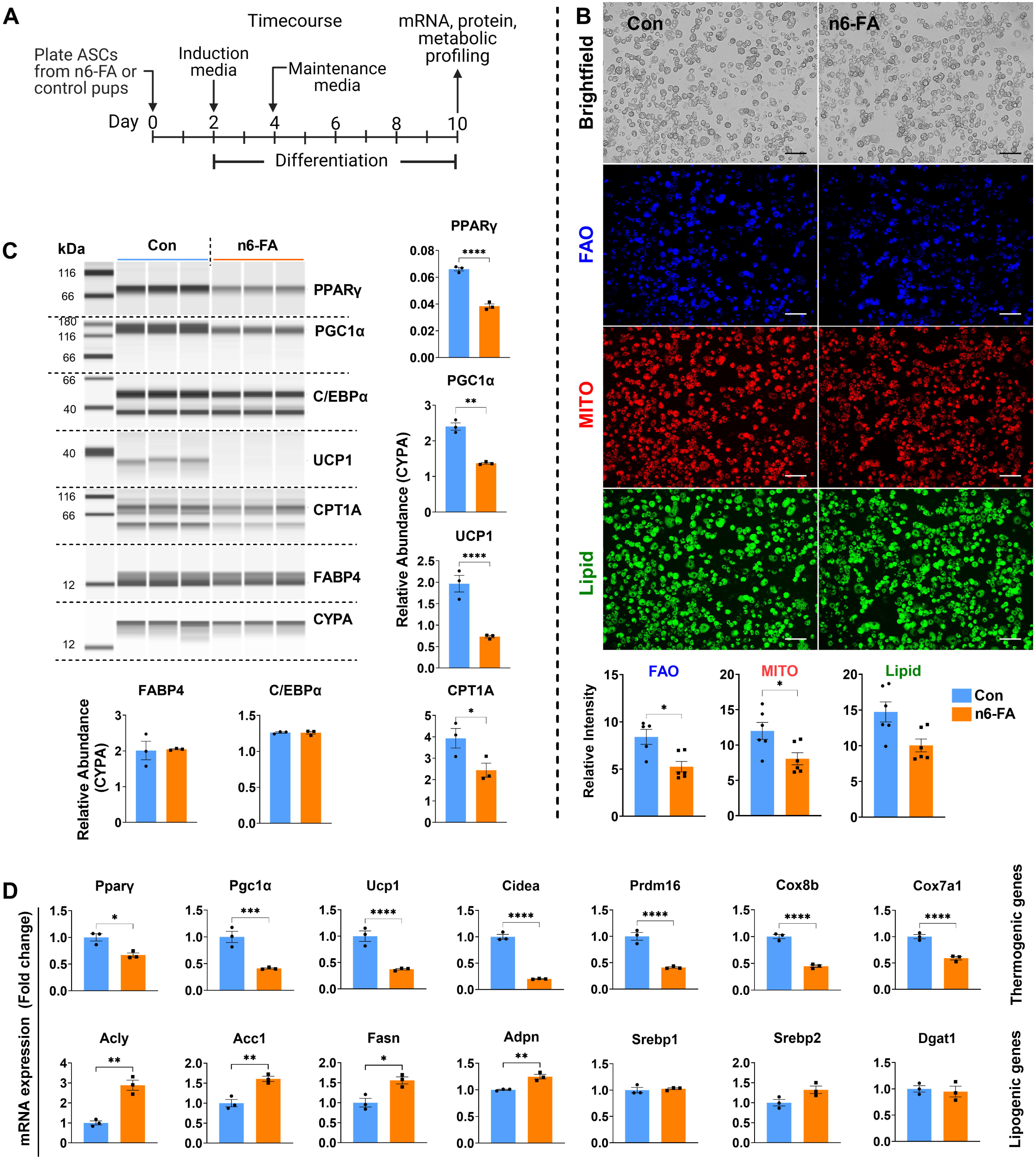
High n6-FA exposure reduces expression of beige markers while upregulating lipogenic markers in adipocytes generated from them. (A) Experimental design. (B) Differentiated beige adipocytes were stained with LipidSpot (green; lipid droplets), TMRE (red; mitochondria), and FAOBlue (blue; fatty acid oxidation). n=3 wells containing ASCs isolated from inguinal SAT from independent litters per control/n6-FA group. Intensity of LipidSpot, TMRE, and FAOBlue staining was quantified using image J software (n=6. Two images per well). Scale bar 100 μM. (C, D) Protein levels and mRNA expression in differentiated adipocytes were quantified through semiquantitative Western blots and qPCR (n=3 wells containing ASCs isolated from inguinal SAT from independent litters per control/n6-FA group). Data are expressed as mean ± SEM, statistical significance is denoted by *p < 0.05, **p < 0.01, ***p < 0.001, ****p < 0.0001 by one-way ANOVA and Tukey’s correction.

### ASCs programmed by n6-FA differentiate into less oxidative adipocytes

Given the decreased FAO and mitochondrial activity measured by live cell stains, we investigated the cellular fuel preferences and oxidative capacities of ASCs isolated from inguinal SAT and differentiated *in vitro*. Adipocytes differentiated from n6-FA ASCs exhibited a marked reduction in FA fuel utilization when administered palmitate, as indicated by the energy map (Fig 4A, red arrow). A sizeable decrease in basal respiration, maximal respiration, ATP production coupled respiration, and spare respiratory capacity was observed in the *in vitro* differentiated adipocytes from n6-FA exposed ASCs, although non-mitochondrial respiration was same for both groups (Fig 4B and C). Furthermore, when treated with mitochondrial FA transporter inhibitor etomoxir, which is used to demonstrate mitochondrial FAO specificity, *in vitro* differentiated adipocytes from n6-FA ASCs were less sensitive (Fig 4D, 20% compared to 55%), suggesting less reliance on FA as mitochondrial fuel. A similar oxygen consumption rate (OCR) profile occurred when glucose was the mitochondrial fuel substrate, with significant decreases basal respiration, and maximal respiration and spare respiratory capacity in the n6-FA group (Fig 4E and F). Addition of the pyruvate carrier inhibitor UK5099 indicated the n6-FA differentiated adipocytes were less sensitive to inhibiting pyruvate entry into mitochondrial oxidative phosphorylation (Fig 4G, 50% relative to 75%). No significant differences were observed when glutamine was the mitochondrial fuel substrate (Fig 4H-J). While nutrient-oxidizing adipocytes are specialized to uptake and oxidize both glucose and fatty acid, nutrient-storing adipocytes divert carbon away from respiration into nutrient storage pathways.

**Figure 4.**
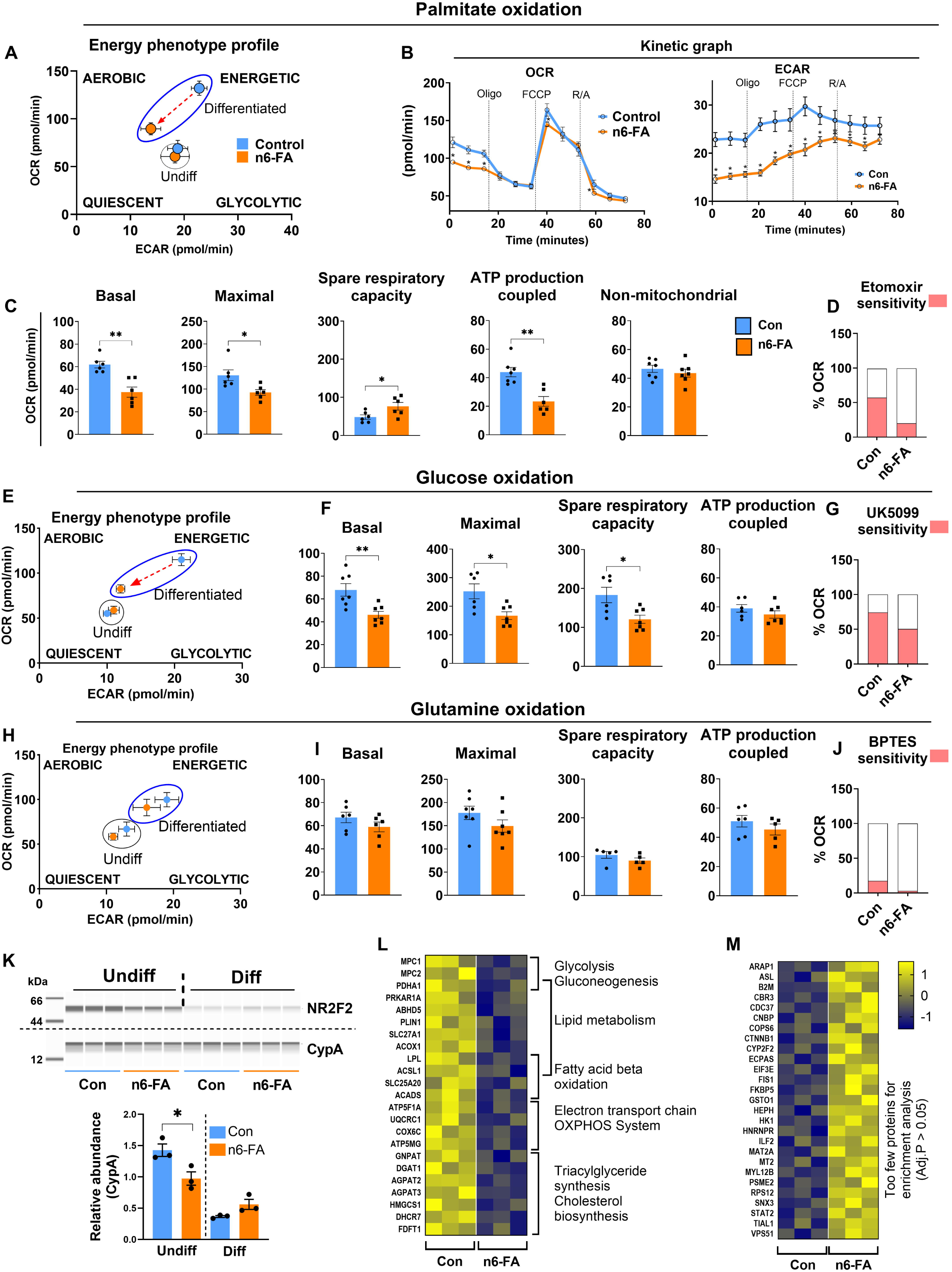
High n6-FA exposure in ASCs inhibits fatty acid oxidation in adipocytes. ASCs were isolated from control and n6-FA exposed pups, plated, and differentiated to beige adipocytes, which were used for metabolic assays in Fig A-J (n=5-7 ASC pools from independent litters per control/n6-FA group). (A) Energy map of control and high n6-FA exposed adipocytes when palmitate was supplied as fuel. (B) Kinetic graphs for oxygen consumption rate (OCR) and extracellular acidification rate (ECAR). (C) OCR for basal, maximal, ATP production-coupled respiration, and spare respiratory capacity in differentiated adipocytes. (D) Sensitivity to Etomoxir during maximal respiration (E, F) Energy map, OCR for basal, maximal, ATP production-coupled respiration, and spare respiratory capacity when glucose was supplied as fuel source. (G) Sensitivity to UK5099 during maximal respiration (H, I) Energy map, OCR for basal, maximal, ATP production-coupled respiration, and spare respiratory capacity when glutamine was supplied as fuel source. (J) Sensitivity to BPTES during maximal respiration (K) Analysis of NR2F2 protein levels in undifferentiated and differentiated beige adipocytes derived from flow-sorted ASCs from control and n6-FA pups through. (L, M) Proteomics data from differentiated beige adipocytes derived from control and n6-FA primary ASCs, followed by pathway enrichment analyses. (n=3 wells containing differentiated ASCs isolated from inguinal SAT from independent litters per control/n6-FA group for all the experiments; K-M). Data are expressed as mean ± SEM, statistical significance is denoted by *p < 0.05, **p < 0.01, ***p < 0.001, ****p < 0.0001 by multiple comparison t-test for B, one-way ANOVA for F, I, K, and t-test for the rest.

### Omega-6 FA ASCs have low NR2F2 levels, giving rise to adipocytes with reduced mitochondrial ETC and FAO enzymes

NR2F2 expression in metabolic tissues has been recognized since 2002^42^. Elegant work by Wang et al. demonstrated that the high-affinity ligand of NR2F2 is an atypical sphingolipid, 1-deoxysphingosine (1-DSO)^37^. We recently reported that NR2F2 is significantly decreased in PND12 pup ASCs following postnatal n6-FA exposure, and that 1-DSO treatment of undifferentiated immortalized ASCs increased beige adipocyte genes prior to differentiation^36^. This finding indicated that activated NR2F2 might alter ASC differentiation to a nutrient-oxidative differentiation trajectory. In isolated primary undifferentiated ASCs, NR2F2 protein was significantly reduced in pups exposed to n6-FA, and following *in vitro* adipocyte differentiation, NR2F2 levels declined equivalently (Fig 4K). In contrast to our previous finding, we differentiated ASCs isolated from control and n6-FA litters into adipocytes for proteomics analyses. Using the significant differentially expressed proteins (DEP), pathway enrichment analysis identified decreased enzymes for lipid metabolism (LPL, PLIN1, ABHD5, SLC27A1), fatty acid oxidation (ACADS, ACSL1, SLC25A20), mitochondrial electron transport chain, and oxidative phosphorylation (ATP5F1A, UQCRC1. COX6C, ATP5MG), glucose metabolism (Mitochondrial Pyruvate Carrier1/2), and cholesterol biosynthesis in n6-FA adipocytes (Fig 4L, Source data file 1). Too few proteins were upregulated to return enriched pathways with an adjusted p-value less than 0.05 (Fig 4M).

### Transient NR2F2 activation before differentiation enhanced FAO after differentiation

Given that NR2F2 protein levels decline following in vitro differentiation, we reasoned that ligand activation of NR2F2 prior to differentiation would confer its function. We treated immortalized ASCs transiently with a 1-DSO pulse (300nM) flanking adipocyte differentiation induction, followed by maintenance media without 1-DSO for six days (Fig 5A). Activating NR2F2 in immortalized ASCs with transient 1-DSO enhanced the energetic profile and significantly increased basal, maximal OCR, and spare respiratory capacity compared to untreated controls (Fig 5B, C). Importantly, transient NR2F2 activation before adipocyte differentiation led to increased beige adipocyte regulators PPARγ, PGC1α and CPT1A after differentiation (Fig 5D). Using primary ASC isolated from n6-FA exposed pups, transient 1-DSO activation of NR2F2 (as in 5A) significantly restored the energetic profile, basal, and maximal FAO and spare respiratory capacity following *in vitro* differentiation into adipocytes (Fig 5E-G). In addition to enhancing FAO and regulators of beige adipogenesis, transient activation of NR2F2 led to persistent protein increases after differentiation for enzymes of mitochondrial ETC, oxidative phosphorylation, sphingolipid metabolism, ATP5 family of ATP synthases, and the NDUF family of mitochondrial Complex I components (Fig 5H, Source data file 1, proteomics, n=3, *pAdj*<0.05). Proteins that were significantly downregulated in response to the transient activation of NR2F2 in n6-FA ASCs included modulators of glucose metabolism, the TCA cycle, and lipid metabolism, including the de novo lipogenic enzymes ACLY, ACACA, ACACB, and FASN (Fig 5I). Together, these findings indicate lasting changes to cellular metabolism resulting from transient NR2F2 activation prior to differentiation in the n6-FA ASCs, ultimately leading to broad changes in the mitochondrial enzymes and function.

**Figure 5.**
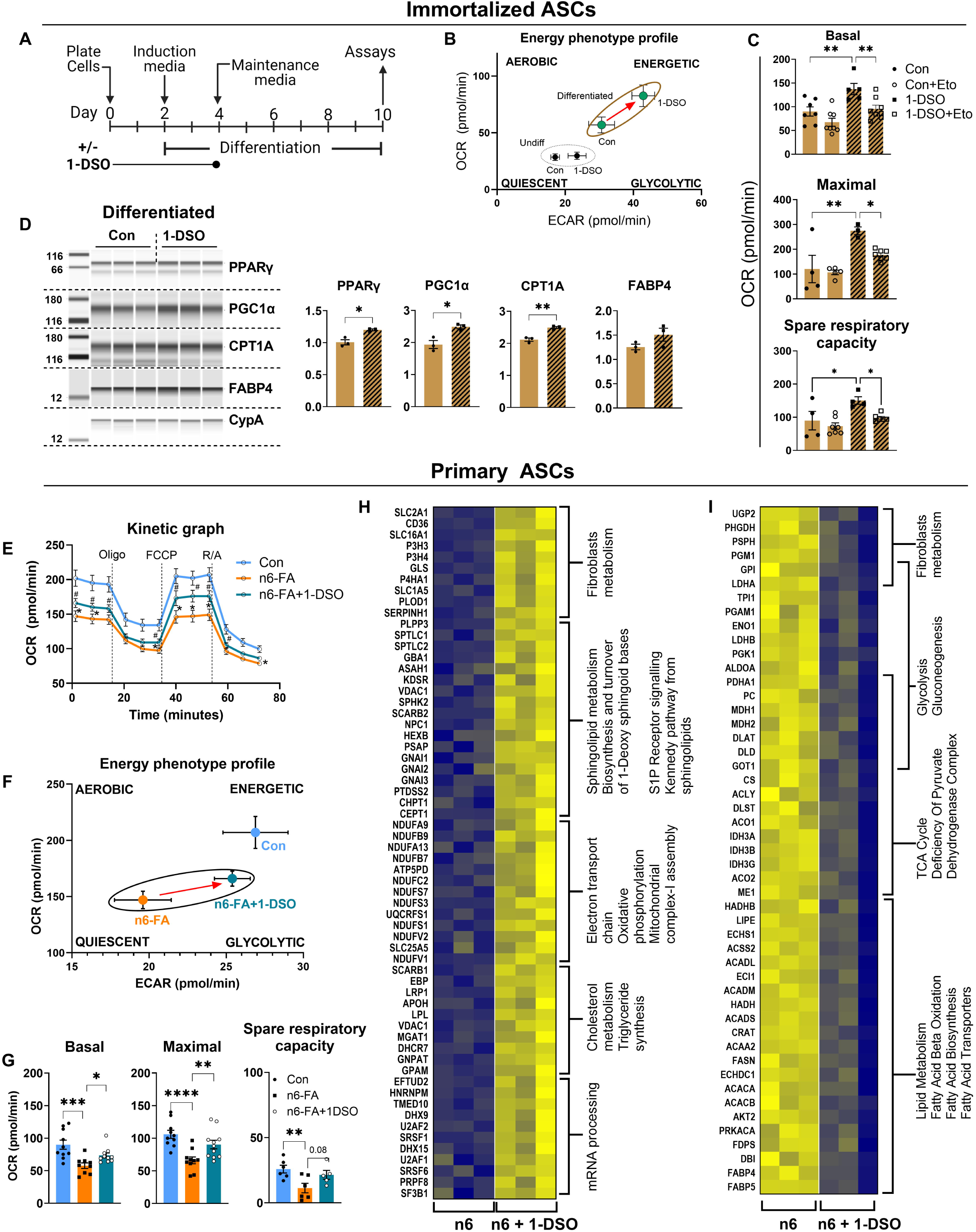
Transient NR2F2 activation in n6-FA ASCs improves fatty acid oxidation and alters mitochondrial proteins in adipocytes. (A) Experimental design. Immortalized adipocyte precursor cells (APCs) were differentiated to beige adipocytes ± a 4-day pulse treatment with 1-DSO in the early phase of differentiation, as shown in the schematic. Following differentiation, cells were assayed for metabolic activity and protein abundance (n=3-7 wells containing immortalized ASCs for Con and 1-DSO; Fig A-D). (B, C) Energy map, basal, maximal respiration and spare respiratory capacity in control and 1-DSO treated beige adipocytes when palmitate was used as the fuel source. (D) Treatment with NR2F2 ligand 1-DSO increased expression of beige markers in beige adipocytes differentiated from APCs evaluated through Western blotting. (E-G) OCR kinetic graph, energy map, basal, maximal respiration and spare respiratory capacity in beige differentiated ASCs (± 1-DSO during differentiation as in panel A) isolated from control and n6-FA pups, when palmitate was supplied as fuel source (n=5-11 wells per group). (H, I) Proteomics data from beige differentiated ASCs (± 1-DSO during differentiation as in panel A) isolated from n6-FA pups, followed by pathway enrichment analyses, showing important metabolic pathways were altered significantly between groups (n=3 wells per group). Data are expressed as mean ± SEM, statistical significance is denoted by *p < 0.05, **p < 0.01, ***p < 0.001, ****p < 0.0001 by one-way ANOVA for C, G, multiple comparison t-test for E and t-test for the rest.

### NR2F2 ablation before differentiation phenocopies the n6-FA nutrient storing adipocytes

Rescue of the diminished FAO in n6-FA exposed pup ASCs by transient NR2F2 ligand activation suggests that NR2F2 operates upstream of, or parallel to, the beige differentiation program during adipogenesis. To test the loss of NR2F2 function and specificity of 1-DSO activation, we isolated ASCs from PND12 inguinal SAT (as above) using NR2F2^f/f^ litters from the control-FA condition and deleted NR2F2 *in vitro* using Adeno-Cre (ΔNR2F2 ASCs) (Fig 6A). These ASCs grew and developed as wildtype ASCs under the *in vivo* control-FA exposure that produces BeAT, robust induction of BeAT regulators, and high levels of mitochondrial metabolism (Figures 2-4). After viral recovery and one passage of the ASCs, they were differentiated into adipocytes (as above), and differentiation potential, protein levels, and cellular metabolism were assessed. NR2F2 loss in ASCs resulted in significantly reduced FAO and mitochondrial activity by FAOBlue and TMRE live cell staining, while lipid droplet accumulation in adipocytes remained unchanged (Fig 6B). Importantly, immunoblotting confirmed that NR2F2 was efficiently deleted following Adeno-Cre transduction in ASCs, and following differentiation, that induction of BeAT regulator proteins, PGC1α and PPARγ, were reduced similarly to the n6-FA exposed inguinal ASCs (Fig 6C). *In vitro* deletion of NR2F2 also resulted in reduction of gene expression for Pparγ, Pgc1α, Cidea, Prdm16, Ucp1, and beige adipogenic regulator Ebf2 (Fig 6D). Deletion of NR2F2 prior to differentiation significantly reduced the OCR using palmitate as the fuel substrate, and transient treatment with 1-DSO failed to restore FAO (Fig 6E,F). These findings demonstrate the specificity of 1-DSO in activating NR2F2 in ASCs and support that NR2F2 is needed for FAO in the n6-FA adipocytes (Fig 5E-F).

**Figure 6.**
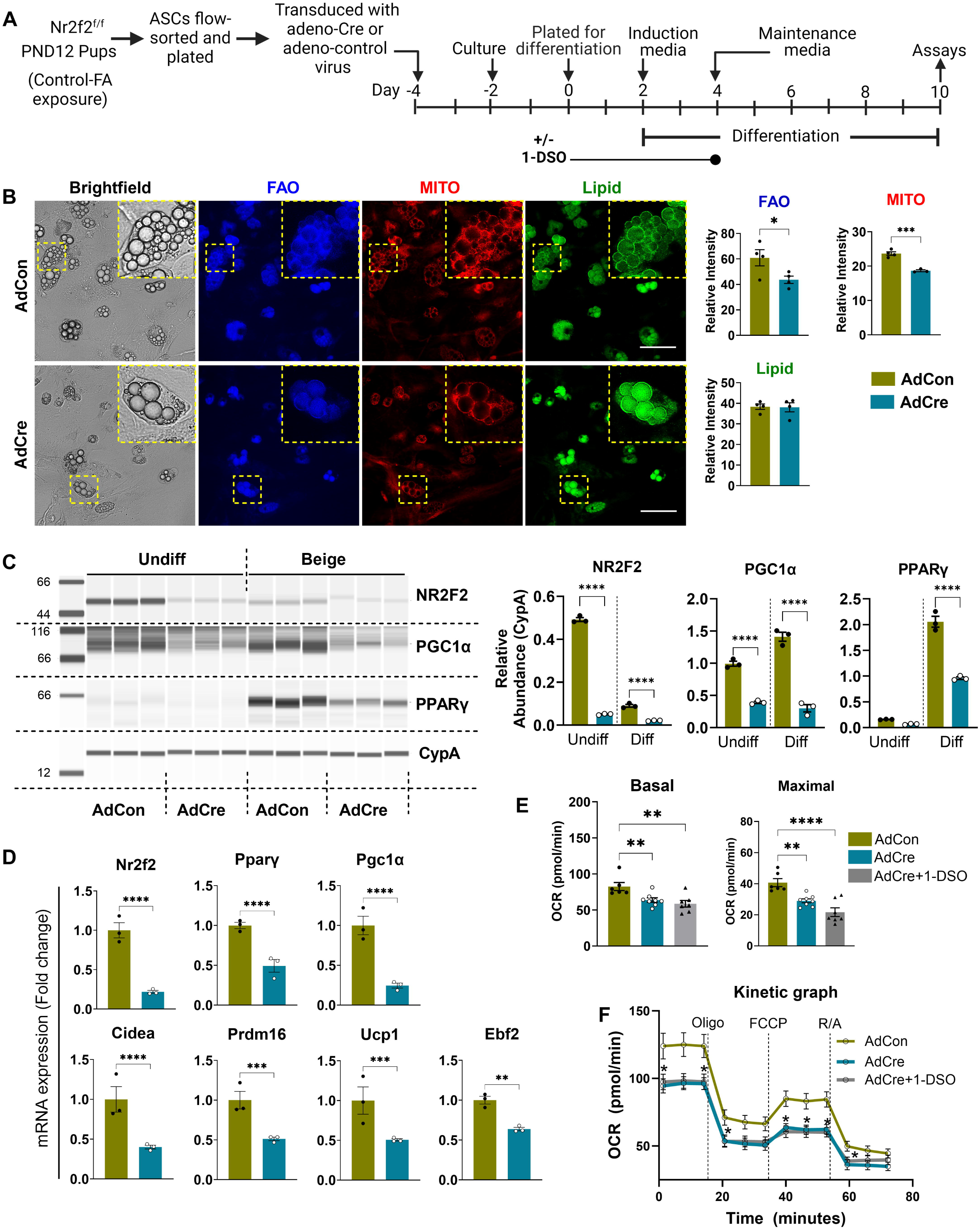
Ablation of NR2F2 *in vitro* suppresses expression of beige proteins following differentiation. (A) Diagram of experimental design. ASCs were isolated from Nr2f2^f/f^ pups, plated, treated with Adeno-Cre (AdCre) or control adenovirus (AdCon) to ablate NR2F2, followed by differentiation into beige adipocytes. Following differentiation, cells were assayed for adipogenic markers. (B) Differentiated adipocytes were stained with LipidSpot (green; lipid droplets), TMRE (red; mitochondria), and FAOBlue (blue; fatty acid oxidation) and quantified in Image J (n=4 wells containing ASCs isolated from inguinal SAT from independent litters). (C, D) Protein and mRNA levels of NR2F2 and other key regulators were quantified in ASCs ± *in vitro* ablation of NR2F2 through Western blotting and qPCR (n=3 wells per group). (E, F) Basal, maximal respiration, and OCR (kinetic graph) of beige differentiated ASCs (n=6-8 wells per group). Data are expressed as mean ± SEM, statistical significance is denoted by *p < 0.05, **p < 0.01 by t-test for B and one-way ANOVA for the rest.

We evaluated the activation status of the Wnt-CTNNb1 pathway using ASCs from the n6-FA and control-FA exposed pups for the known Wnt-Ctnnb1 responsive gene Axin2 ^57^. In undifferentiated ASCs, Axin2 gene expression was significantly decreased, indicating that Wnt-CTNNb1 pathway was less active in n6-FA ASCs compared to the ASCs isolated from the control-FA pups. Continuous stimulation of the WNT/CTNNb1 signaling pathway in the 3T3L1 preadipocyte cell line led to chronic overexpression of NR2F2^38^. In alignment with the NR2F2 protein levels (Fig 4K), gene expression level of Nr2f2 was significantly decreased in the n6-FA ASCs, and mRNA for the Sptlc2 enzyme responsible for synthesizing 1-DSO was not significantly different (Fig 7A). Importantly, in n6-FA ASCs administered a selective CTNNb1 stabilizer to mimic WNT signal transduction activation of CTNNb1, the expression of Nr2f2, Sptlc2, and Axin2 were induced in ASCs isolated from the n6-FA exposed pups. Taken together, this indicates that CTNNb1 activation in primary ASCs induces Nr2f2 in the n6-FA exposed ASCs, linking WNT-CTNNb1 signal transduction to NR2F2 induction in primary ASCs.

**Figure 7.**
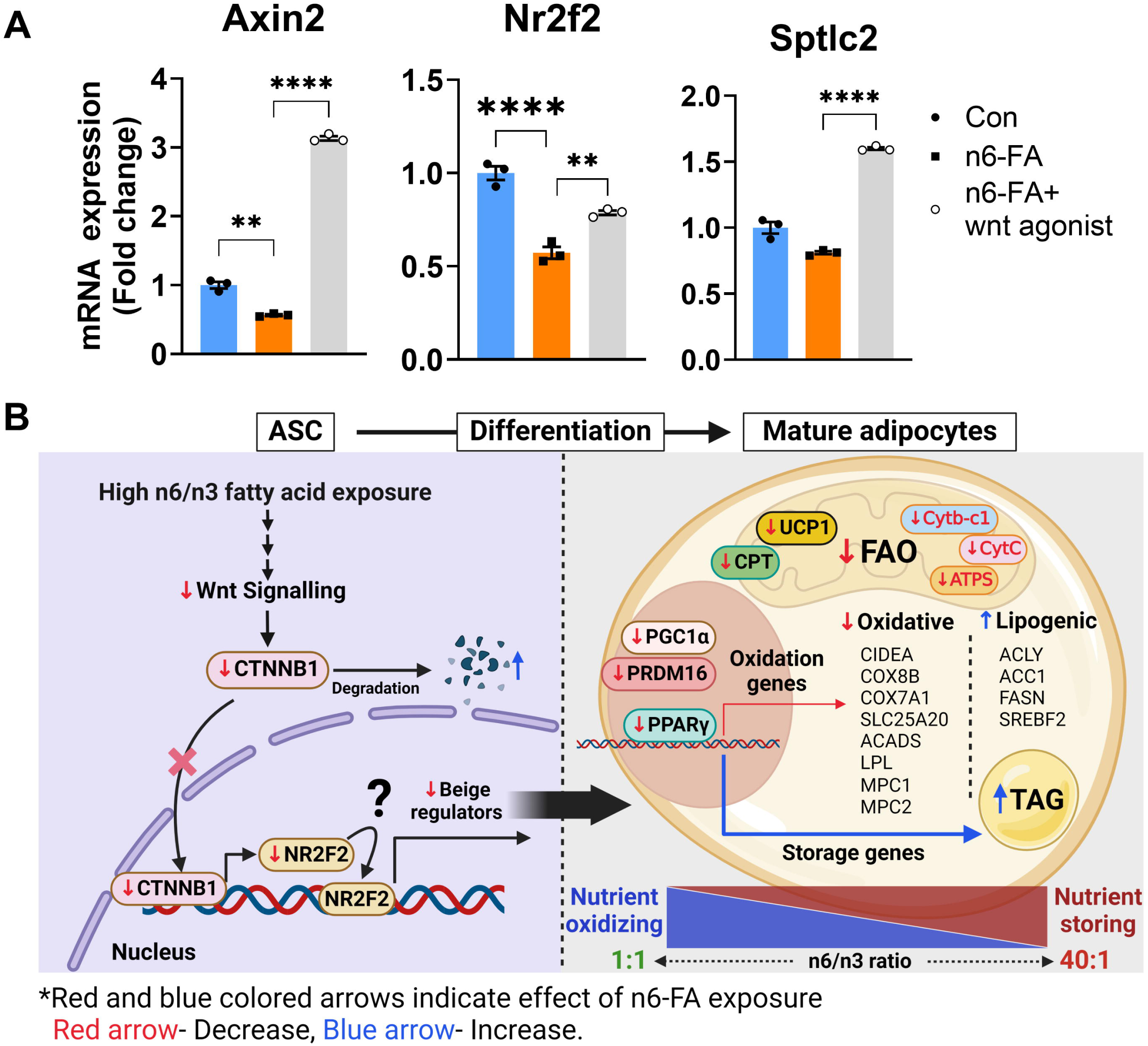
(A) mRNA expression of Nr2f2, Axin2, and Sptlc2 in ASCs ± Wnt agonist 1(48h) quantified through qPCR (n=3 wells per group). (B) Model showing effects of excess n6-FA exposure on ASCs and how that affects adipocyte metabolism. ASCs exposed to n6-FA ratio have a diminished WNT mediated activation of CTNNb1 gene signature, leading to poor induction of NR2F2 and lower expression of beige regulators PPARγ, PGC1α, Cidea, and Prdm16. These ASCs differentiate, give rise to mature adipocytes with diminished electron transport chain and oxidative phosphorylation protein components, blunted nutrient oxidation, while retaining an enhanced capacity for triacylglyceride accumulation. Data are expressed as mean ± SEM, statistical significance is denoted by *p < 0.05, **p < 0.01, ***p < 0.001, ****p < 0.0001 by one-way ANOVA.

## DISCUSSION

Lipids transmitted from mother to infant provide energy dense nutrients, structural components, and potent signaling molecules for adipogenesis^58^. Over the first six months, infants significantly increase their body fat^59^. Early-life body fat accumulation serves dual roles, nutrient storage and oxidation, the latter of which is needed to defend core body temperature and supply energy needed for growth^60,61^. Both n6- and n3-FA are essential early in life for brain and eye development, but they also shape the development of the fat depots, possibly setting the stage for future obesity risk^62,63^. We investigated whole body and cellular energetics of PND12 pups using a model of excessive perinatal n6-FA exposure, at a point in life when metabolically active BeAT is highly abundant^64^. Litters exposed pre- and postnatally to a disproportionately high n6/n3 FA ratio during adipose development had a higher RER, reduced lipid oxidation calculated by the equation 1.70 × VO_2_ – 1.69 × VCO_2_, and diminished whole-body FAO by tracing ^13^C-palmitate oxidation at 26°C (Fig 1C-E). The diminished FAO in litters occurred in the face of equivalent palmitate levels present in both milk from dams (i.e., FA intake) and in circulating plasma of the pups (Table1), suggesting that n6-FA exposed litters had sufficient palmitate to oxidize but did not utilize it as a preferred fuel source. Analysis of the ^13^C-palmitate uptake indicated no differences for metabolic tissues, including BAT, liver, inguinal SAT, and muscle (Supp Fig 1B). Together, these findings suggest that the high n6-FA exposure changed the fuel utilization preference away from lipid oxidation, because the capacity to oxidize lipid as a fuel source was blunted by exposure to high levels of n6-FA *in vivo*. A shift away from lipid fuel preference is consistent with the metabolic programming we previously reported in the adult setting, where pups exposed to a high n6/n3 FA ratio during the perinatal window, had significantly higher RER and predisposition for adipose accumulation in adulthood^30^.

This metabolic phenotype characterized by diminished whole-body FAO is, at least in part, due to a defective metabolism within the SAT, given that n6-FA exposed SAT had a white adipocyte morphology, size distribution, and 40% increase in stored triacylglycerides (Fig 1F-H). The SAT morphology and lower whole-body FAO is consistent with the n6-FA pups having less overall BeAT *in vivo.* The molecular signature of isolated adipocytes from PND12 n6-FA exposed pups supports this observation, with less PGC1α, PPARγ, CPT1A, CPT1C, and UCP1, which are key regulators of adipose mitochondrial FAO. PPARγ is necessary for both beige and white adipogenesis, and zinc finger protein 423 (ZFP423) is thought to act as a molecular “switch” between these two developmental programs^65^. In the beige program, EBF2 binds beige gene promoters and recruits co-activators, which mediate decondensation of chromatin structure and recruitment of PPARγ, promoting expression of beige loci and therefore beige adipogenesis. In the white program, ZFP423 binds to EBF2, recruiting the NuRD corepressor complex to block EBF2-dependent chromatin decondensation and subsequent PPARγ recruitment, thereby promoting the nutrient storing white adipocyte program and not the nutrient oxidizing beige one^65^. Conditional ablation of ZFP423 in inguinal white adipocytes increased levels of Pgc1α, Prdm16, and Ucp1 expression, leading to a beige adipocyte morphology in the presence of the potent PPARγ agonist rosiglitazone^65^. Intriguingly, NR2F2 has recently been shown in a chicken model of adipogenesis to repress expression of ZFP423 by binding to sites in the ZFP423 promoter, toggling to the beige adipocyte program^66^. In this study, we show that NR2F2 levels are reduced in ASCs following perinatal exposure to high levels of n6-FA, associating with a significant reduction of PGC1α, UCP1, and PPARγ *in vivo* (Fig 2), as well as numerous beige genes (Fig 3) following *in vitro* differentiation. Taken together, our results suggest a model for NR2F2 in which reduction of NR2F2 permits expression of ZFP423, which in turn, blocks EBF2 function, promoting the white adipogenesis program. Consistent with our current understanding of the beige and white adipogenesis is the loss of beige gene expression and reciprocal induction of lipogenic genes, including Acly, Acc1, Fasn, and the principal regulator of the cholesterol biosynthesis pathway, Srebf2 (Fig 3).

In alignment with the diminished whole-body FAO observed by ^13^C-palmitate tracing was the marked decrease in cellular FAO of *in vitro* differentiated ASCs isolated from the n6-FA exposed pups (Fig 4). When provided palmitate as the fuel source, adipocytes differentiated *in vitro* had a 50% reduction in basal, maximal, and ATP-coupled respiration, with less sensitivity to the mitochondrial FA uptake inhibitor etomoxir. Interestingly, when glucose was provided as the cellular metabolic fuel substrate, both basal and maximal respiration rates were also reduced, while minimal differences were observed with glutamine (Fig 4). Together, this finding suggests a fundamental difference in the oxidative phosphorylation capacity of adipocyte mitochondria following a high n6-FA perinatal exposure. The pathway enrichment analyses of the proteome from *in vitro* differentiated ASCs supports this notion, in that, downregulation of OXPHOS and ETC proteins ATP5F1A, ATP5MG, ACOX, ACADS, COX6C, and UQCRC1 was observed (Fig 4).This indicates a significant difference in the key enzymes responsible for mitochondrial oxidation and energy production, which is a well-documented characteristic of mitochondrial dysfunction of adipocytes in obesity^67,68^, which is established as early as PND12 in mice. The observations of impaired *in vitro* palmitate and glucose substrate oxidation, the loss of beige adipocyte regulator levels (Fig 3), and 40% more triacylglyceride in the PND12 fat pad (Fig 1), is consistent with adipogenic programming that is prone to store nutrients rather than oxidize them.

We identified previously that NR2F2 was reduced in isolated inguinal fat pad ASCs from n6-FA exposed pups, alongside an altered mitochondrial gene expression profile^36^. We extend those findings to show that ligand activation of NR2F2 in ASCs leads to persistent formation of nutrient oxidizing adipocytes. NR2F2 is expressed during development and is critical for energy homeostasis and adipocyte biology^39–43^. As a nuclear receptor type 2 family member, NR2F2 can bind DNA directly, as well as dimerize with other nuclear receptors to either activate or repress transcription, depending on the cellular context^69^. While NR2F2 is known as a metabolic regulator, defining the mechanisms of NR2F2 function *in vivo* has remained a challenge. There is conflicting evidence of NR2F2 function during adipogenesis^39–44^. Genomic loss of Nr2f2 is embryonic lethal in mice^70^, and the genomic heterozygous mice are viable but smaller in size than WT littermates^71^, have more skeletal muscle, less white AT, and greater bone formation^39,40^. Paradoxically, the genomic Nr2f2 heterozygous adult mice had greater energy expenditure, glucose clearance, and resistance to diet-induced obesity, thought to be due to the imbalance of metabolic tissues ^39,40,42,69,71^.

Metabolically active adipocytes are established by known regulatory factors, most prominently PPARγ, PGC1α, and PRDM16, which cooperate to implement mitochondrial gene expression as part of the beige adipocyte metabolic program^55,72^. Here, the perinatal n6-FA exposure led to isolated ASCs with significantly lower NR2F2 protein levels (Fig 4K). We found that transient NR2F2 activation in ASCs led to persistent increases of metabolic activators PPARγ, PGC1α, and CPT1A, along with partially restoring the diminished FAO capacity due to perinatal n6-FA programming (Fig 5). Although NR2F2 activation with 1-DSO was transient, the whole cell proteome revealed installation of lasting increases in mitochondrial complex I, oxidative phosphorylation, and electron transport chain components (Fig5 H, I).

The increase in mitochondrial complex components supporting FAO corresponded to decreases in proteins for glycolysis, TCA cycle, *de novo* fatty acid synthesis, and curiously, some enzymes for FAO and fatty acid transport. In other words, NR2F2 activation increased levels of some enzymes responsible for FAO, while simultaneously decreasing enzymes involved in the same process. Perhaps this dual effect resulted in adipocyte mitochondria with an increased ability to handle fatty acids nearly as effectively as adipocytes differentiated from control ASCs. Importantly, 1-DSO treatment is specific to NR2F2 activation in ASCs, because transient treatment failed to boost FAO following *in vitro* ablation of NR2F2 (Fig 6). Instead, ASCs isolated from NR2F2^f/f^ pups, which developed normally as wildtype ASCs under the control-FA perinatal exposure, accumulated lipid droplets equivalently to AdCon-transduced control ASCs, but lacked robust induction of Pparγ, Pgc1α, Cidea, Prdm16, and Ucp1 beige genes when NR2F2 was knocked down. Altogether, ASC-specific decreases of NR2F2 that occur during perinatal n6-FA exposure coincided with diminished FAO, which could be partially restored by activating NR2F2 with ligand 1-DSO in the undifferentiated state.

It remains unclear what regulates the levels of NR2F2 mRNA and protein in ASCs from our model of perinatal FA exposures. In the 3T3-L1 preadipocyte cell line, chronic WNT3a-stimulated activation of CTNNb1 in undifferentiated cells and throughout 10-days of adipocyte differentiation, induced and sustained NR2F2 levels in adipocytes, suppressing adipogenesis via modifications of chromatin architecture^38^. Because NR2F2 normally decreases during adipocyte differentiation (Fig4K), sustaining its levels throughout differentiation may have influenced those findings. Despite the evidence that CTNNb1 activation opposes adipogenesis, key effectors of WNT signaling (CTNNb1 and TCF7L2) are expressed in mature adipocytes, and disrupting them leads to adipocyte hypertrophy, inflammation, glucose intolerance, and insulin resistance^73–75^. Recent findings highlight the potential of “WNT positive ASCs” in promoting beige adipocyte formation and recruitment, and when transplanted, WNT positive ASCs enhanced glucose metabolism in recipients^76^. We previously reported that ASCs from n6-FA pups expressed an mRNA signature consistent with inhibited CTNNb1, including downregulated Wingless-type MMTV integration site 6 and 9a (Wnt6 and Wnt9a), its receptor Frizzled 1 (Fzd1), and signal transducer Akt1, which have been shown to activate CTNNb1 ^76^. Conversely, n6-FA exposed ASCs had upregulated secreted Frizzled Related Protein 2 and 4 (Sfrp2 and Sfrp4), which have been shown to antagonize the WNT/CTNNb1 signaling pathway^77^.

Interestingly, n3-FA supplementation has been shown to increase Wnt/CTNNb1 levels and signaling *in vitro* and in animal models. For example, pregnant female rats that were induced to have preeclampsia had increased Wnt and b-catenin protein levels in their brain tissue when they were supplemented with n3-FA using both eicosapentaenoic and docosahexaenoic acids blended at 4:1^78^. In another study, supplementation of docosahexaenoic acid in doses above 10mM increased WNT/CTNNb1 signaling in human iPSC-derived neuronal progenitor cells *in vitro* in the presence of Wnt ligand (Wnt3a)^79^. In our study, ASCs from n6-FA exposed pups had significantly downregulated Axin2, a known CTNNb1 target gene. Stimulation of n6-FA ASCs with a selective CTNNb1 stabilizer significantly increased Axin2, which coincided with induced gene expression of Nr2f2 and Sptlc2, an enzyme responsible for synthesizing 1-DSO^37^, the endogenous ligand for NR2F2 (Fig 7A). Upregulation of Nr2f2 mRNA following CTNNb1 activation in the n6-FA primary ASCs is consistent with NR2F2 being downstream of WNT/CTNNb1 gene regulation.

Cumulatively, our findings support a model in which ASCs exposed to a high n6-relative to n3-FA ratio tapers WNT mediated activation of CTNNb1, leading to reduced induction of NR2F2 and less robust expression of beige regulators PPARγ, PGC1α, Cidea, Prdm16, and Ucp1. These ASCs, when differentiated, give rise to mature adipocytes with diminished electron transport chain and oxidative phosphorylation protein components, blunted nutrient oxidation, while retaining enhanced triacylglyceride accumulation (Fig 7B). Our findings suggest that, from as early as PND12 in mice, inguinal SAT may be programmed for nutrient storing white adipocytes, potentially leading to an obesity-prone phenotype that persists into adulthood, possibly via epigenetic regulation^30^.

## LIMITATIONS AND FUTURE RESEARCH

This study differed from our previous studies of early-life FA exposures and SAT development ^30,36^, in that maternal dietary FA composition was based on extremely n6-FA rich safflower oil. Although dams and sires were provided the safflower oil diet at the time of pairing, thereby confining the FA exposure within the perinatal setting, we cannot exclude the possibility of germline effects^80^. Moreover, this dietary FA composition is not representative of the US maternal dietary n6-relative to n3-FA ratio, which has escalated dramatically over the decades due to an overload of n6-FA^11^, reaching an n6/n3 ratio between 20:1 and 40:1^81^. Because dietary oil sources, such as corn, soybean, sunflower oils, are typically high in n6-FA, an important issue will be to lower maternal consumption of n6-FA containing sources while increasing dietary sources containing n3-FA, such as flax and fish oils^62^. The current study did not investigate the classes of signaling lipids, which are enzymatic products of both n6- and n3-FA. For example, our group recently identified 12,13-diHOME, an oxylipin derivative of linoleic acid (18:2 n6) that is inversely related to infant fat mass at 1-month of age^82^. Interestingly, NR2F2 activating ligand 1-DSO was not detected in either the dam’s milk or pup’s plasma by lipid mass spectrometry, suggesting that it is synthesized as an intracellular ligand or acts as a localized paracrine signaling lipid. To that end, it would be important to evaluate the intracellular signaling lipids, including the NR2F2 ligand 1-DSO, within ASCs isolated from n6-FA and control-FA exposures, for differences in lipids influencing adipogenic potential. Critically, NR2F2 binds CTNNb1^83^, suggesting that direct CTNNb1-NR2F2 interaction may play a key role in modulating how ASCs respond during adipogenesis early in life. Although we suggest that NR2F2 acts upstream of the beige regulator genetic program, it is unclear whether NR2F2 is upstream of, or acting in parallel to, these other important regulators. Future work will focus on defining the sequential timing of NR2F2 action and any ASC-specific NR2F2 interacting proteins that might differ between control and n6-FA perinatal exposures.

## CONCLUSION

Obesity, defined by excessive white adipose tissue accumulation, is a complex gene-nutrient-lifestyle interaction disease, with roots in early-life development. The metabolic regulator NR2F2 plays a key role in establishing adipocytes with the capacity to oxidize nutrients, at least in the mouse pup. While still somewhat controversial in humans, activating beige adipocytes in rodents offsets metabolic dysfunction associated with diet-induced obesity. Understanding mechanisms governing how metabolically active adipogenesis is established, especially in response to dietary bioactive FA and their signaling lipid derivatives, could pave the way for promising interventions that protect against childhood obesity.

## ACKNOWLEDGEMENT

We would like to thank the Flow cytometry core at The University of Oklahoma health Sciences center (OUHSC). We are also thankful to the Flow cytometry core at Oklahoma Medical Research Foundation for assistance with Aurora Spectral Flow Cytometry (Grant number: 1S10OD028479-01). This work is supported by grants: R24GM137786 (IDeA National Resource for Quantitative Proteomics) and P20GM103447 (Oklahoma INBRE) to MK, NIH (HL 156856, HL 137799) and AHA (TPA97002) to PRN, and Oklahoma Center for Adult Stem Cell Research (OCASCR) and the Presbyterian Health Foundation Equipment Grant to MCR.

## AUTHOR CONTRIBUTION

**Conceptualization-** MCR, SD. **Methodology-**SD, RV, GPM, AEM, JWF, MK. **Investigation-**MCR, SD, RV, GPM. **Data Curation-** SD, RV, AEM, GPM, GKD, KH, MK, JWF. **Formal analysis-** MCR, SD, RV, JWF, MK. **Resources**-MCR, PRN. **Writing-Original Draft-** SD, MCR, GPM, RV. **Writing-Review and Editing**-MCR, GPM, RV, PRN. **Visualization**-MCR. **Supervision**-MCR. **Funding acquisition**-MCR.

## ETHICS DECLARATIONS

### Competing Interests

The authors declare no competing interests.

## INCLUSION AND DIVERSITY

One or more of the authors of this paper self-identifies as an underrepresented ethnic minority in their field of research.

## Notes

### Competing Interest Statement

The authors have declared no competing interest.

