## Supplemental Figure 1 for "NR2F2 Reactivation in Early-life Adipocyte Stem-like Cells Rescues Adipocyte Mitochondrial Oxidation"

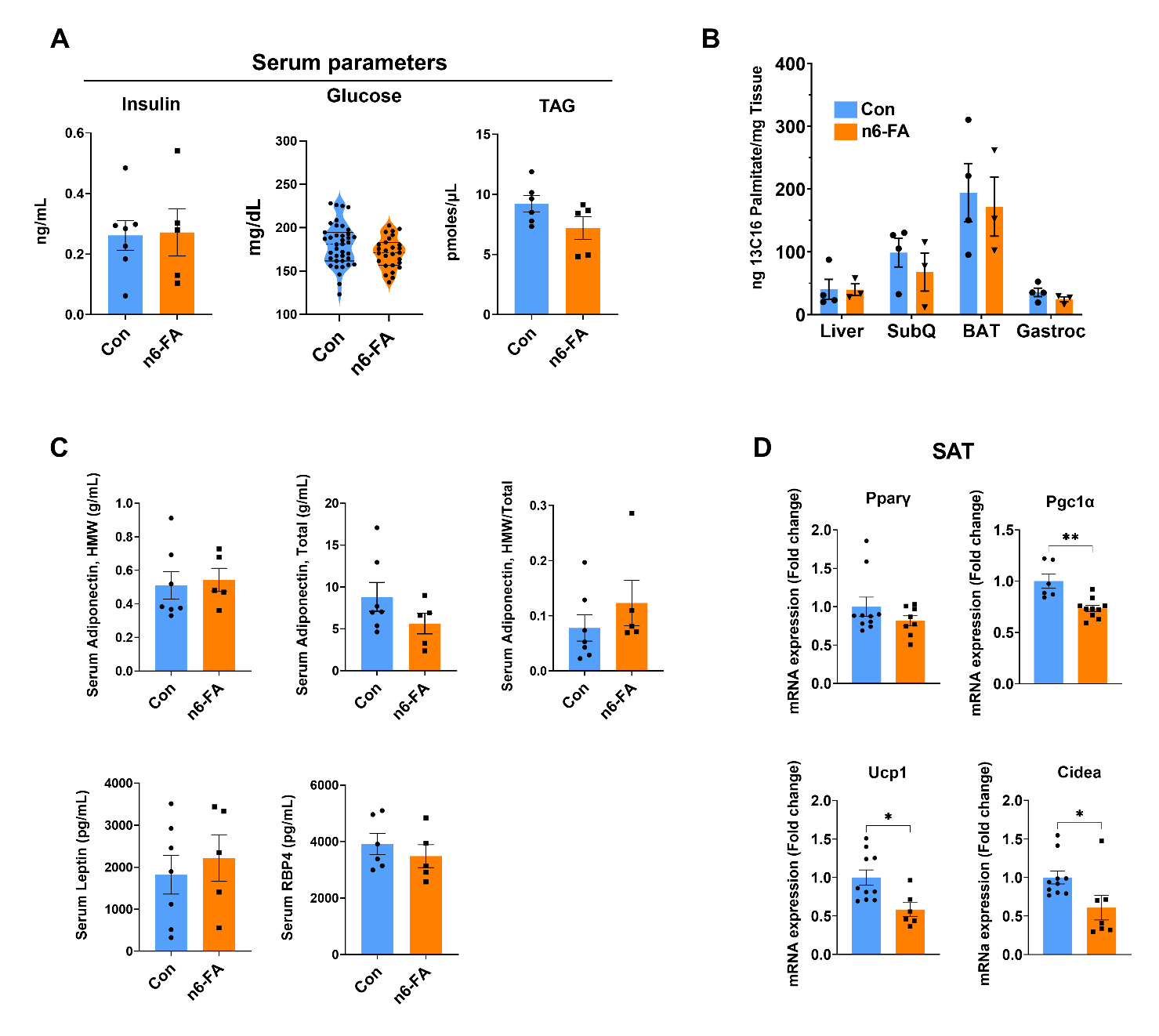
 **Supplemental Figure 1.** (A) Blood insulin, glucose, and TAG levels in control and n6-FA exposed pups (n=5-7 for Insulin and TAG and 27-39 for Glucose). (B) ^13^C_16_-palmitate uptake measured by GC/MS analysis of liver, inguinal subcutaneous fat, BAT, and gastrocnemius muscle (n=3-4). (C) Serum levels of adiponectin (total and high molecular weight), leptin, and RBP4 quantified by ELSA (n=5-7). (D) mRNA expression of Pparγ, Pgc1α, Ucp-1, and Cidea in inguinal fat pad measured through qPCR (n=6-10). Data are expressed as mean ± SEM, statistical significance is denoted by *p < 0.05, **p < 0.01, ***p < 0.001, ****p < 0.0001 by t-test for A, C, D, and one-way ANOVA for B.
