## Supplemental Figure 2 for "NR2F2 Reactivation in Early-life Adipocyte Stem-like Cells Rescues Adipocyte Mitochondrial Oxidation"

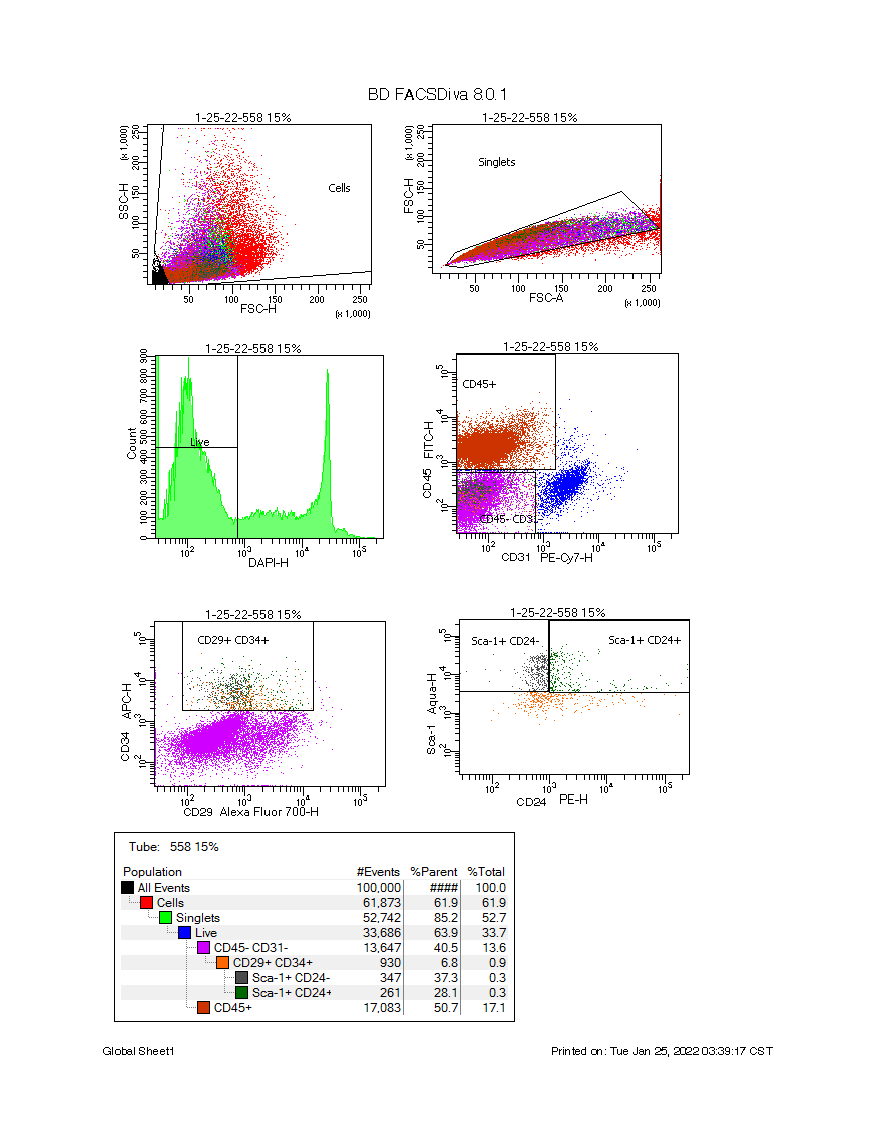
 **Supplemental Figure 2**. Gating strategy to obtain ASCs from stromal vascular fraction of subcutaneous white adipose tissues from a litter of PND12 pups.
